# Assessing the ligand native-like pose using a quantum mechanical-derived hydropathic score for protein-ligand complementarity

**DOI:** 10.1101/2025.10.20.683430

**Authors:** Brian Medel, William J. Zamora, Enric Herrero, Glen E. Kellogg, Jana Selent, Javier Vázquez, F. Javier Luque

## Abstract

Elucidating the correct binding mode of drug-like compounds is crucial to disclose the molecular determinants that underline the recognition by the target protein and estimate the binding affinity, thus guiding the ensuing hit-to-lead optimization. However, the choice of the near-native binding pose in docking experiments remains a major hurdle. Assuming that the hydropathic complementarity principle is the major driving force in ligand-protein recognition, the suitability of simple hydropathicity-based scoring functions to reveal the near-native pose was examined. A benchmarking dataset of 1000 ligand-protein complexes purposely designed to encompass bioactive and decoy poses was used to assess the performance of 3D hydropathicity atomic descriptors derived from empirical and quantum mechanical models. This strategy led to a predictive accuracy of ca. 90% when the non-negligible influence of the conformational stress is also considered. The results reveal the need to leverage nonpolar/polar chemical features for the successful discrimination of the near-native pose.

## Introduction

Elucidating the bioactive pose of a ligand is essential for understanding the molecular determinants that underlie the recognition and binding between a small molecule and the target protein.^1–3^ This knowledge is fundamental for the success of a drug design campaign; thus, predicting the correct binding mode of a drug-like candidate allows medicinal chemists to suggest suitable chemical changes based on the specific 3D geometrical and physicochemical features of the ligand-protein complex in the hit-to-lead optimization. However, the identification of the bioactive pose from the chemical structure of the drug-like compound remains one of the greatest challenges in computer-aided drug design, especially when high-throughput screening methods must face chemical libraries containing millions to even billions of compounds.^4–8^

Docking-based methods are the most popular strategy to explore the feasibility of small molecules to interact with the target protein, providing information about the 3D arrangement at the binding cavity, often with scores that aim to reflect the complementarity between the chemical features of both ligand and pocket residues. The performance of docking methods in terms of speed primarily depends on the efficiency of the geometrical searching algorithms used to generate poses compatible with the size and shape of the pocket, while the discrimination between the ligand poses in the binding cavity is dictated by tailored scoring functions.^9–11^ Discriminating between poses is challenging due to the limited accuracy of the scoring functions, which *a priori* must condense the distinct enthalpic and entropic contributions in a single score. Over the years a variety of strategies have been formulated, which can be categorized into physics-based, empirical and more recently machine-learning-based scoring functions. ^12–16^

Among the many properties influencing ligand binding, desolvation is one of the major components, and hence the hydropathic complementarity between ligand and receptor is widely assumed to be a key requirement for the binding affinity. Indeed, the importance of both the geometrical features of the pocket and hydropathicity has been highlighted in several studies aimed at predicting target druggability and the binding affinity.^17–21^ Specifically, ligand desolvation is largely responsible of the variation in maximal achievable binding energy for a drug-like molecule.^22–24^ In this context, it is not surprising that previous efforts have attempted to develop strategies exploring the 3D hydropathic pattern of ligands, as noted in the heuristic molecular lipophilic potential, which relies on the analysis of the electrostatic potential at the molecular surface to provide a unified lipophilicity and hydrophilicity potential, and the Hydropathic INTeraction (HINT) scoring function, which exploits atomic contributions to the *n*-octanol/water partition coefficient (LogP) from experimentally measured values of small chemical moieties to quantify the hydrophobic interactions between ligand and protein.^25,26^ This simple, yet effective approach has proven to be successful in many drug discovery campaigns. ^27–30^

Building on HINT strengths, our aim is to explore the feasibility of a physics-based hydropathic scoring function to determine the bioactive pose of a protein-ligand complex. Instead of depending on empirical LogP libraries, our approach is focused on quantum mechanical (QM) calculations of the solvation free energy of ligands in *n*-octanol and water, coupled to a perturbative scheme that permits decomposition of the molecular LogP into atomic contributions, leading to the previously described HyPhar descriptors.^31–33^ Noteworthy, this strategy considers the specific chemical features (protonation, tautomerism, conformation) of the ligand’s bioactive species. Furthermore, this approach also includes the changes in enthalpy and entropy that underscore the hydropathic complementarity required for the mutual recognition between ligand and protein residues.

In this work, we examine the suitability of the 3D hydropathicity atomic descriptors derived from empirical and QM-derived models in conjunction with a simple mathematical expression to discriminate between near-native and decoy poses of protein– ligand complexes. To achieve this, we created a publicly available dataset that includes native and five decoy poses for 1000 ligand-protein complexes chosen to represent the content of PDBBind. ^34^ This dataset allows evaluation of how well the hydropathic complementarity principle alone can identify the bioactive or near-native pose of the ligands. As a final test, we assess the performance of the scoring function using a collection of G protein-coupled receptor (GPCR) complexes, which encompass one of the most relevant pharmaceutical families in drug discovery.

## Results

### Database of ligand-protein complexes

To evaluate the performance of the hydropathic scoring function for discriminating amongst ligand poses bound to the target protein, we generated a dataset, named DB1000, comprising 1000 ligand-protein systems selected from the complexes present in PDBbind 2020. DB1000 reflects the distribution of functional families as indexed in the Protein Data Bank (Fig. 1A).^35^ The physicochemical properties of the ligands in DB1000 encompass a diversity chemical space, ranging from fragment-like compounds to drug-like candidates (Fig. 1B). Thus, their molecular weights vary from *ca*. 150 to 600 Da, containing compounds with 2-8 rotatable angles. While most of the ligands have no formal charge, there is a significant number of positively/negatively charged molecules. Similarly, almost 60% are achiral and *ca*. 30% contain one or two chiral centers. Finally, most of the compounds contains between 1 and 4 hydrogen-bond donors, and between 2 and 9 hydrogen-bond acceptors. Overall, whereas these features resemble the general trends of the Lipinski’s rule-of-five, they reflect the chemical diversity of the ligands.

**Figure 1.**
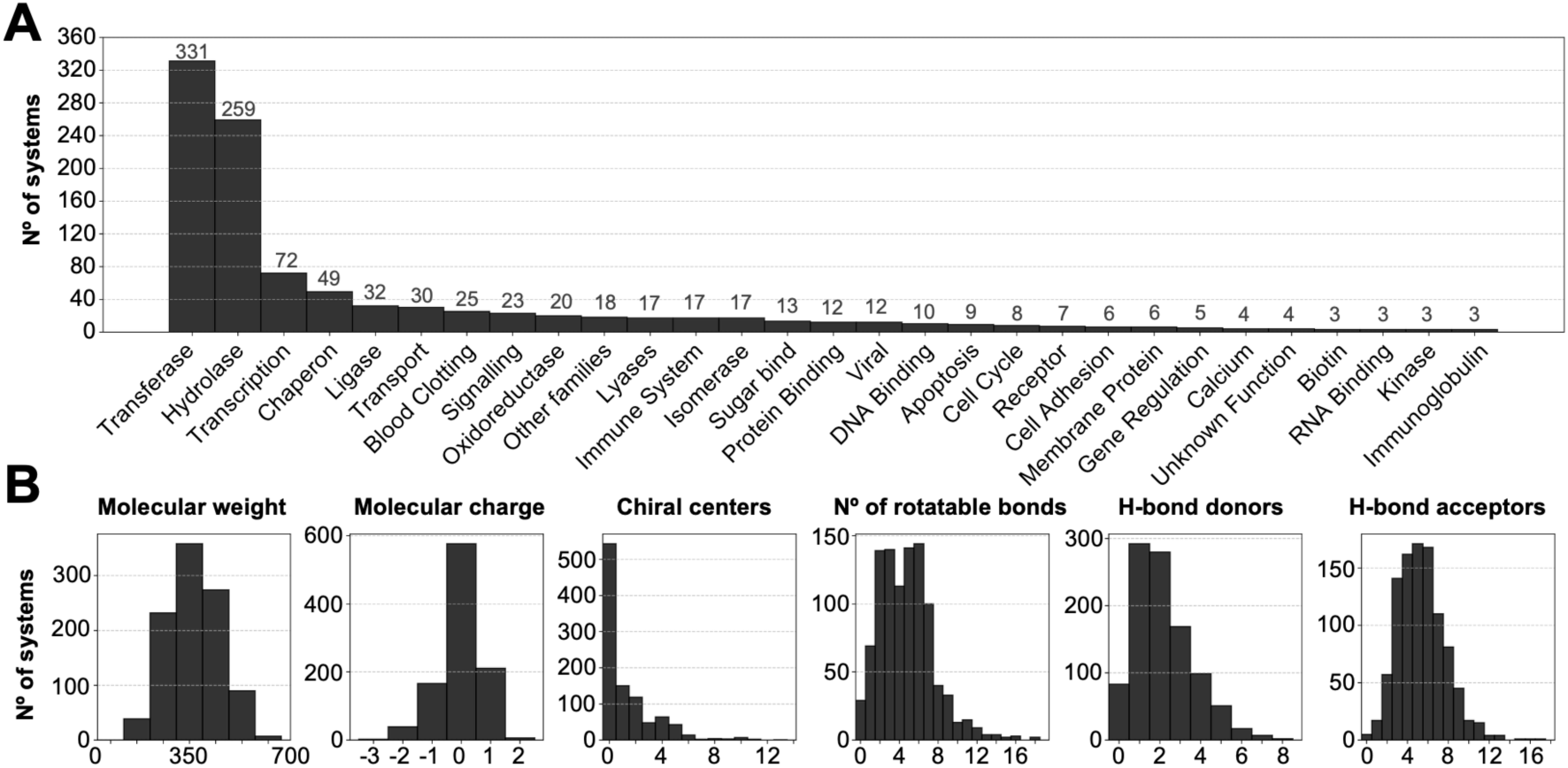
Classification of proteins in the DB1000 dataset and analysis of the ligand chemical space. (A) Distribution of ligand-protein complexes in DB1000. Only families with more than three representatives are shown, all others are grouped under the “Other” category. The main families encompass 33% (transferases) and 26% (hydrolases) of the complexes in the database. (B) Population distribution of molecular weight and formal net charge, number of chiral centers, number of rotatable bonds, and number of hydrogen-bond donors and acceptors for the set of ligands included in DB1000.

After preparation of each ligand-protein complex (Fig. 2A; see also Methods), docking calculations were performed to generate five decoy poses of the ligand using Glide. ^36^ Accordingly, in addition to the crystallographic pose, each target protein is associated to five ligand poses suitably chosen with positional root mean-square deviations (RMSD) relative to the heavy atoms of the X-ray pose in the ranges <2, 2–4, 4–6, 6–8 and >8Å (e.g., Fig. 2B).

**Figure 2.**
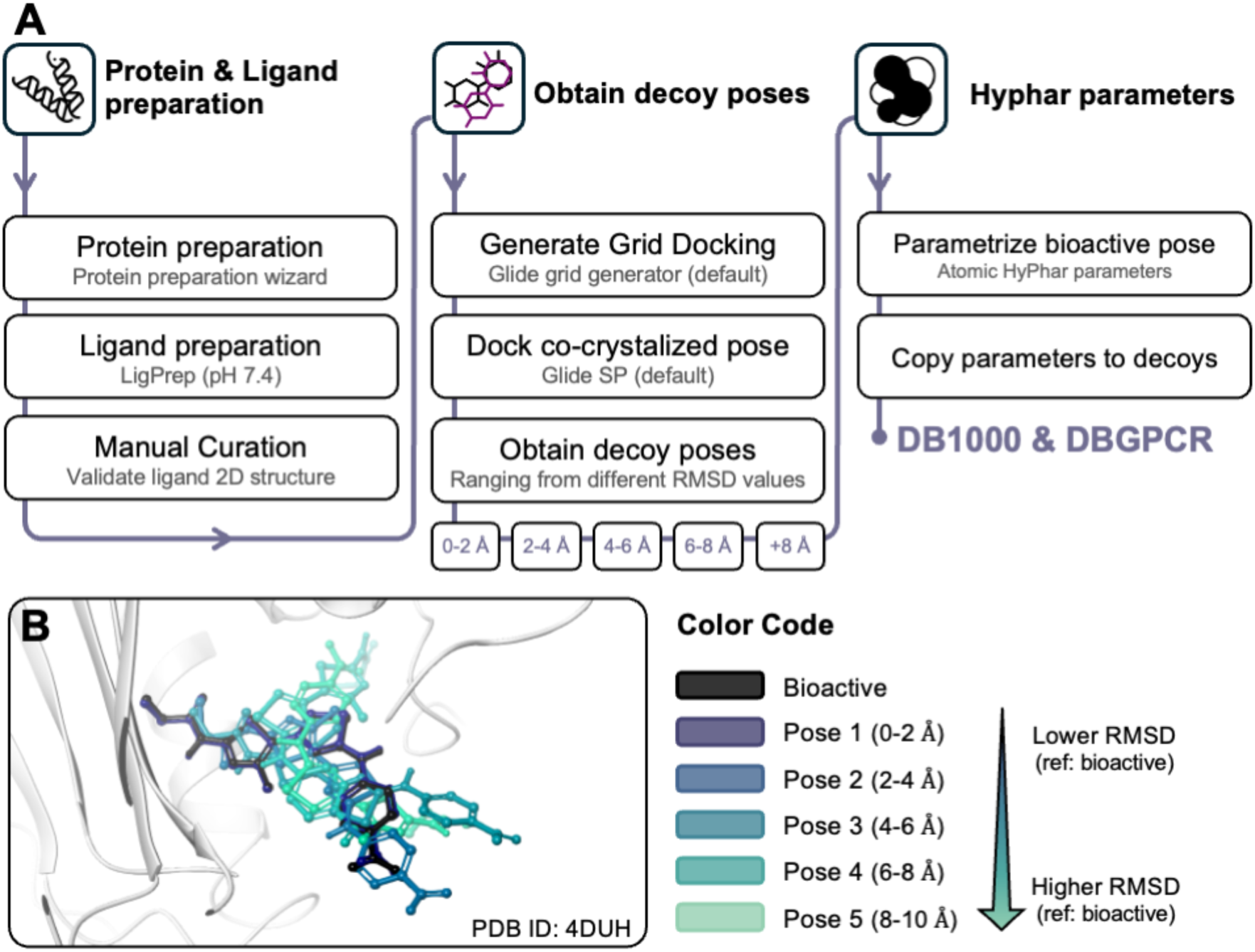
Generation of ligand poses in DB1000. **(A)** Schematic pipeline used to generate the benchmark datasets. **(B)** Color-coded representation of the ligand poses generated to assess the discriminating ability of the hydropathic scoring function to identify the bioactive pose exemplified for the complex of 4-{[4’-methyl-2’-(propanoylamino)-4,5’-bi-1,3-thiazol-2-yl]amino}benzoic acid with *E. coli* DNA gyrase B (PDB ID 4DUH). The X-ray pose is depicted in black, and the docked poses, varying from low RMSD to high RMSD, transition from dark blue to light green.

### The hydropathic scoring function

To examine the reliability of the hydropathic complementarity to discriminate between the X-ray or near-native binding mode from decoy poses of the ligand, we have implemented a hydropathic scoring function model (*S*_*hydro*_; Eq. 1) inspired by the HINT model of Kellogg and Abraham:

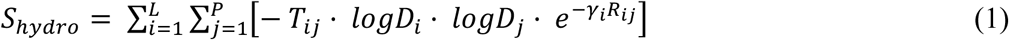

where *i* and *j* are indexes for the atoms in the ligand and protein, respectively, *logP*_*i*_ and *logP*_*j*_ represent the hydropathic contributions of atoms *i* and *j* in the ligand and the protein, respectively, *T*_*i*j_ is an integer term (1 or -1) that controls the pairwise attractive or repulsive behaviour of the pairwise interaction between atoms, specifically necessary for polar atoms that may be acids or bases, and the exponential function *e*^-*γiRi*j^ accounts for the distance-dependent interaction between atoms *i* and *j*, which are separated at a distance *R*_*i*j_, and *γ*_*i*_ modulates the decay of the interaction between atoms (see Methods).

The atomic hydropathic contributions of the ligand (HyPhar descriptors) were determined from QM continuum solvation calculations using the B3LYP/6-31G(d) parametrized version of the IEFPCM/MST solvation model (see Methods).^37,38^ The pH-dependent protein-like lipophilicity scale reported in previous studies for the natural amino acids, which was determined at the same level of theory, was used to describe the atomic contributions of the residues in the target pocket.^39^ Accordingly, a consistent computational framework was adopted to quantitatively estimate the hydropathic complementarity for the ligand-protein complexes.

The accuracy of *S*_*hydro*_score was compared with the HINT score (*S*_*HINT*_; Eq. 2)^25^, which evaluates the atomic pairwise interaction between the hydropathic contribution obtained using a partition scheme of experimentally derived LogP of small fragments, based on the CLOG-P scheme of Hansch and Leo.^40^

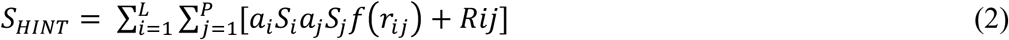

where *a*_*i*_ and *a*_*j*_ denote the hydropathic atomic contributions of ligand and protein, respectively, weighted by the solvent-exposed surface area of the interacting atoms (*S*_*i*_ and *S*_*j*_), *f*(*r*_*i*j_) is an exponential functional form to account for the separation between interacting atoms, and *Rij* is an optional implementation of the Lennard-Jones potential.

The results show that *S*_*hydro*_identifies the bioactive (X-ray) or the native-like (pose 1) correctly in 84.0% (74.3% considering only the docked poses 1-6) of ligand-protein complexes (Fig. 3A). This compares with 60.1% of cases correctly identified by *S*_*HINT*_(Fig. 3A). Pose 2 (2 < RMSD < 4Å) encompass 9.3% of the predictions made by *S*_*hydro*_. They generally correspond to cases showing significant overlap with the X-ray pose within the innermost pockets of the binding cavity, and the structural difference arises from chemical moieties mainly located in solvent-exposed areas, as noted for the ligands in PDB IDs 4QD6, 4KYH, 4I9U and 5ULA, thus facilitating the adoption of distinct arrangements (e.g., flip of cyclic moieties) for the solvent-exposed fragment of the compound (Fig. 3C). In a few cases, the ligand was buried in the pocket, but the topological features of the cavity permitted the adoption of distinct conformations of elongated chains through rotatable bonds, such as PDB ID 2YIX, or the interchange of structurally similar fragments in the ligand, such as PDB ID 4MC6 (Fig 3D). Finally, values less than 3.0% for more deviated decoy poses. This behaviour is not reflected in the *S*_*HINT*_ results, as decoy poses 2-5 are more evenly scored.

**Figure 3.**
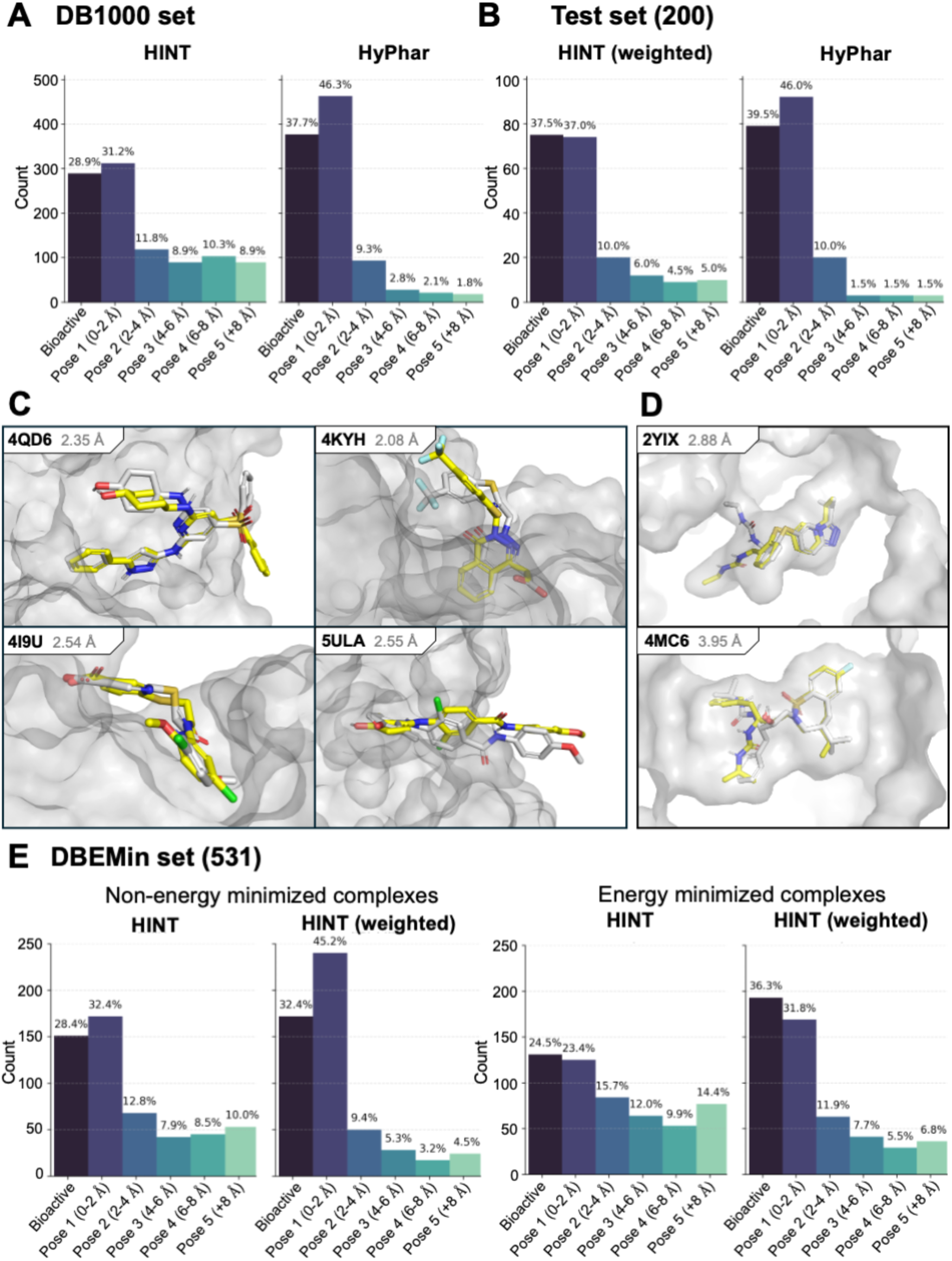
Performance of the scoring functions*S*_*hydro*_ *and S*_*HINT*_. **(A)** Distribution of the top-scored poses obtained for the hydropathic scoring functions *S*_*HINT*_ and *S*_*hydro*_for the complexes in DB1000. **(B)** Distribution of the top-scored poses obtained for the hydropathic scoring functions *S*_*HINT*_ and *S*_*hydro*_ for the test set (200 complexes). **(C, D)** Representative cases of the top-scored pose 2 is the top-scored reflecting changes between the X-ray (gray sticks) and pose 2 (yellow sticks) in (**C**) solvent-exposed terminal regions of the molecule or **(D)** similar arrangements of flexible regions of the ligand in buried cavities. **(E)** Effect of the geometrical refinement of the ligand-protein complexes on the performance of *S*_*HINT*_ for 531 targets included in the DBEMin dataset with and without inclusion of weights for the pairwise interaction components of HINT (left and right plots denote the distribution of poses obtained with and without energy-minimization).

The *S*_*HINT*_score distinguishes six pairwise atomic interactions: three attractive terms, which correspond to the formation of hydrogen bond, acid–base (e.g., salt bridge), and hydrophobic contacts, and three repulsive terms, which account for the misfit between ligand and protein due to the formation of acid–acid, base–base, and hydrophobic-polar contacts. Each of these interactions contribute equally to the *S*_*HINT*_ score, which assumes additivity amongst these interaction types. However, this assumption may be inaccurate if the hydrophobic/hydrophilic balance of the whole molecule is affected by the nature of the chemical groups present in a compound and/or the protein’s binding site. In addition, the exponential decay with distance of the pairwise interaction between atoms as coded in HINT, which has some experimental basis for hydrophobic-hydrophobic interactions, may not ideal be for other interaction classes, causing another possible source of non-additivity.^41^ Accordingly, a weighted sum of the six pairwise atomic interactions might be valuable to correct for deviations from additivity inherent in the unoptimized total *S_HINT_*.

To this end, we randomly partitioned the DB1000 dataset into training (800 complexes) and test (200 complexes) subsets. The former was used to determine the weights for the six classes of pairwise interactions using a Bayesian optimization, and the optimized weights were then included in Eq. 2 and applied to the test set. This process was repeated 25 times, leading to average weights of 1.00, 0.58, 0.56 and 0.45 for hydrophobic, hydrogen-bond, acid-base and base-base contacts, and a negligible weight (0.05) for acid-acid and hydrophobic-polar interactions, thereby indicating the lack of relevant misfits in the docked poses (see Supplementary Material Fig. S1). The accuracy of *S*_*HINT*_in discerning the bioactive or native-like poses increased from 60.1% to 74.5%, while also improving the discrimination of significantly deviated decoy poses (Fig. 3B). Despite this significant improvement in performance, the *S*_*hydro*_ scoring function still outperforms the predictive power for the test complexes, as it reaches an accuracy of 85.5% for the test subset (Fig. 3B).

Since HINT was originally created as a tool for evaluation and subsequent optimization of crystallographic structures, its application to high-throughput and rapid virtual screening challenges the fundamental assumptions underlying its original design. HINT is extremely sensitive to structural details, with large score variances often due to a single non-optimal hydrogen bond. To explore the impact of geometry refinement, a subset of complexes was generated by minimizing the six ligand-protein complexes obtained for each protein using the OPLS4 force field (see Methods).^42^ This led to a subset of 531 complexes, denoted DBEMin, that maintained the proper RMSD distribution of the native-like and decoy poses in the range of RMSD values comprised between <2 and >8 Å. The energy minimization resulted in a slight rearrangement of the binding pocket (on average, 0.6 Å as measured from the RMSD to the X-ray crystallographic structure) and had a similar effect for both X-ray and decoy poses (see Supplementary Material Fig. S2). The application of HINT to the energy minimized complexes yielded an accuracy of 47.9%, which increased to 68.1% when the weighted pairwise contributions were considered (Fig. 3E). These values are in contrast with accuracies of 60.8% and 77.6% for the non-energy minimized complexes, reflecting the sensitivity of *S*_*HINT*_ to subtle structural adjustments that may to favor decoy poses. A similar trend was observed for the performance of *S*_*hydro*_, as the sum of bioactive and native-like poses decreased from 84.4% to 73.2% for the energy-minimized complexes (Supplementary Material Fig. S3). This finding is not unexpected keeping in mind that the tuning of the decay parameter (*γ*_*i*_) assigned to the ligand atoms was adjusted using the X-ray complexes without any energy-minimization refinement.

### Tuning of the hydropathic scoring function

Although the discrimination of the X-ray and native-like pose from decoys supports the reliability of *S*_*Hydro*_ to re-rank the poses generated in docking calculations, several factors may underscore the failure of *S*_*Hydro*_in predicting the bioactive pose in several cases. These factors may reflect the omission of the conformational stress in distinct poses of the ligand, which would affect the probability for distinguishing between poses with low- and high internal energy, the degree of ligand burial in the binding pocket for distinct poses, which would affect the desolvation cost, and the formation of distinct interaction patterns between poses, which might introduce inaccuracies into the pairwise formalism of the hydropathic score.

To examine the potential contribution of these factors, we i) determined the conformational energy of the distinct poses using the MMFF94s force field^43^, ii) estimated the desolvation free energy taking advantage of the known atomic contributions to the hydration free energy of the ligand in the bound and unbound states (see Methods), and iii) counted the number of pairwise interactions that a pose forms with residues in the protein cavity for salt bridge (*n*_*SB*_), hydrogen (*n*_*h*–*bonds*_) and halogen (*n*_*hal*_) bonds, aromatic *π*-*π* and T stacking (*n*_*π*–*st*_, *n*_*T*–*st*_), and cation-*π* contacts (*n*_*cat*–*π*_; see Methods). For the sake of comparison, the values determined for the distinct poses of a ligand-protein complex were normalized to a zero mean and unit variance. Finally, a Bayesian optimization was used to determine the weights of the combinations between the *S*_*Hydro*_score with each separate factor (see Methods). It is worth noting that the conformational energy and desolvation cost were shown to not correlate with the hydrophobic/hydrophilic score, whereas a mild correlation was observed between this property and the number of pairwise interactions (see Supplementary Material Fig. S4).

While the weights obtained upon combination with the conformational energy support the leading role of the hydrophobicity/hydrophilicity (*w*_*hydro*_ = 0.76), the conformational penalty may be significant for a proper discrimination between poses (*w*_*conf*_ = 0.24). Indeed, the accuracy in discriminating the bioactive/native-like pose for the 200 complexes included in the DB1000 test set improved from 85.5% (Fig. 3B) to 89.7% (see Supplementary Material Fig. S5). Although this trait may be affected by the limited accuracy of force fields in describing the relative energy between conformations, the slight improvement supports the hydropathic complementarity as an effective criterion for discriminating the correct pose.

On the other hand, comparison of the differences in desolvation energy that may arise from the distinct burial of the poses in the protein cavity had a marginal impact, as noted in an overall accuracy of 85.1%, which reflects the negligible weight of the desolvation contribution (*w*_*solv*_ =0.09). This effect can be largely attributed to the similar burial exhibited by the X-ray and decoy poses in most ligand-protein complexes (see Supplementary Material S6). Similar trends were observed for the number of protein-ligand contacts formed by the distinct poses, since the accuracy was 86.7%, and the weight of the component was 0.1. This likely reflects the redundancy in the information provided by this variable with the hydropathic complementarity, and the main role of hydrophobicity in guiding the binding affinity of ligands.

### Analysis of the hydropathic profile of binding pockets

The previous results point out that a precise definition of the hydropathy of ligands is key for assessing the complementary overlap with the nonpolar/polar patches of the binding cavity, and hence for discriminating the bioactive pose from decoys. Indeed, the accuracy of *S*_*Hydro*_ depends on the nature of the protein family (Fig. 4A), and this dependence may qualitatively be justified considering the balance between nonpolar and polar patches in the binding cavity. In this regard, the accuracy in discriminating the bioactive pose tends to increase as the number of nonpolar patches decreases, and concomitantly as the ratio of polar atoms increases (Fig. 4B). This suggests that the ability to elucidate the bioactive pose will be challenging for exceedingly large nonpolar, ‘greasy’ cavities, whereas a minimum number of polar groups distributed onto the cavity surface is required to effectively anchor the ligand. On the other hand, the hydropathic profile of the binding pocket is more relevant than the overall size of the pocket, which exhibits a less pronounced influence of the accuracy of *S*_*Hydro*_ (Fig. 4B).

**Figure 4.**
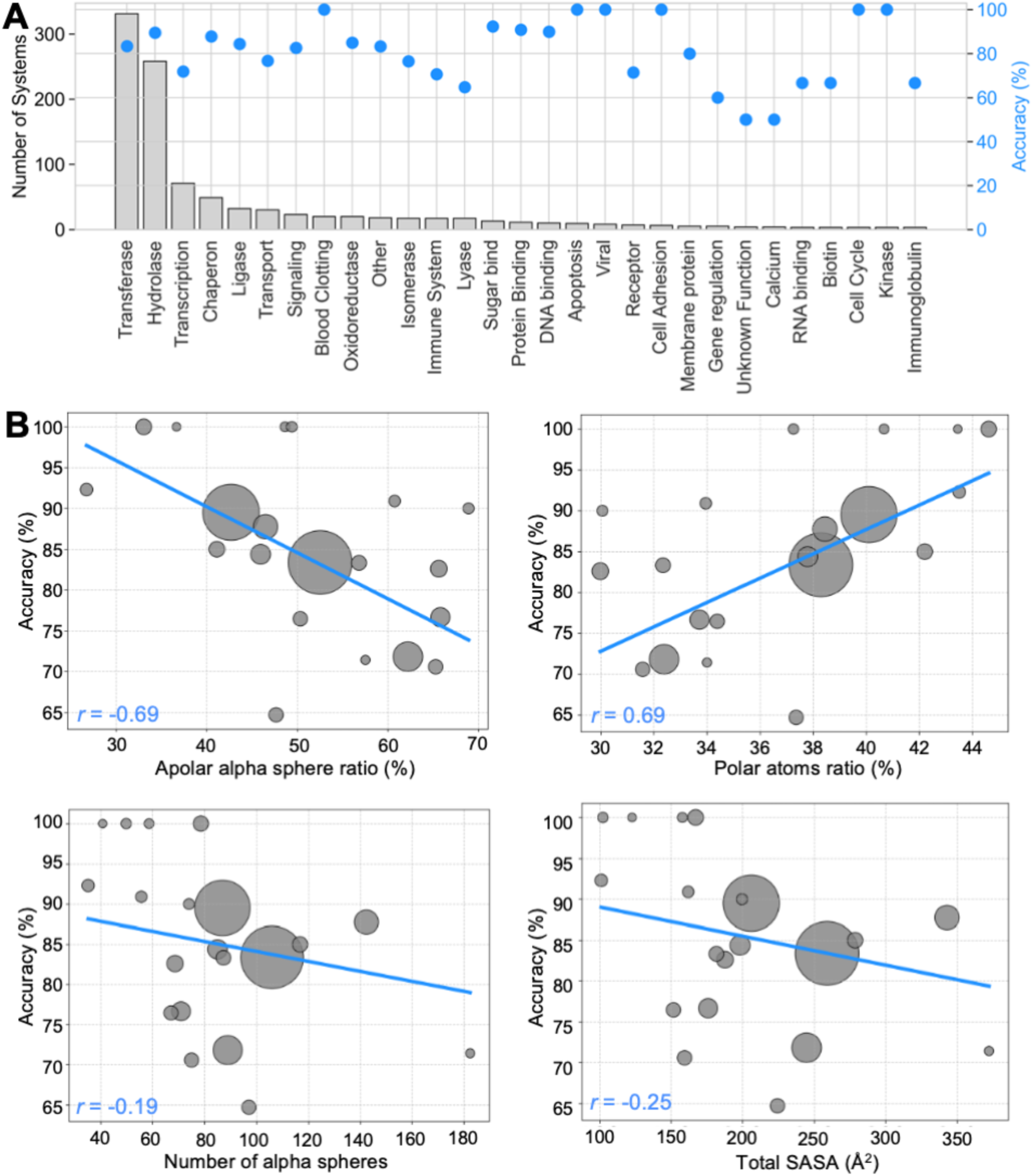
Effect of the nonpolar/polar balance in the binding pocket on the accuracy of *S*_*hydro*_. **(A)** Accuracy per family distribution in the DB1000 dataset using HyPhar descriptors. Total population (bars) and accuracy (blue dots) of each family are shown in left and right y axes. **(B)** (Top) Representation of the accuracy determined for the 200 complexes included in the DB1000 test set versus (left) the ratio of nonpolar patches (measured from the number of nonpolar alpha spheres in the binding cavity) and (right) the ratio of polar atoms present in the cavity walls. (Bottom) Representation of the accuracy determined for the 200 complexes included in the DB1000 test set versus the volume of the cavity (measured from the number of alpha spheres in the binding pocket) and the solvent-accessible surface area of the pocket. All the descriptors were estimated using Fpocket.^44^ Complexes with less than five members are not shown for the sake of clarity.

To check the suitability of the previous findings, we have examined the reliability of the hydropathic complementarity principle to discriminate the native-like poses of a set of ligands present in 196 complexes involving G-protein coupled receptor (GPCR) proteins (denoted hereafter as the DBGPCR database; see Methods), which is used as an external test set. Using the same procedure outlined above to generate decoy poses, the direct application of *S*_*Hydro*_ leads to an accuracy of 67.4%, which would increase up to 80.3% upon inclusion of the poses with a RMSD between 2 and 4 Å (Fig. 5A). Noteworthy, if one limits the analysis to the docked poses, excluding the X-ray arrangement of the ligands, the performance of *S*_*Hydro*_ compares with the ranking provided by the Glide SP scoring function (Fig. 5B).

**Figure 5.**
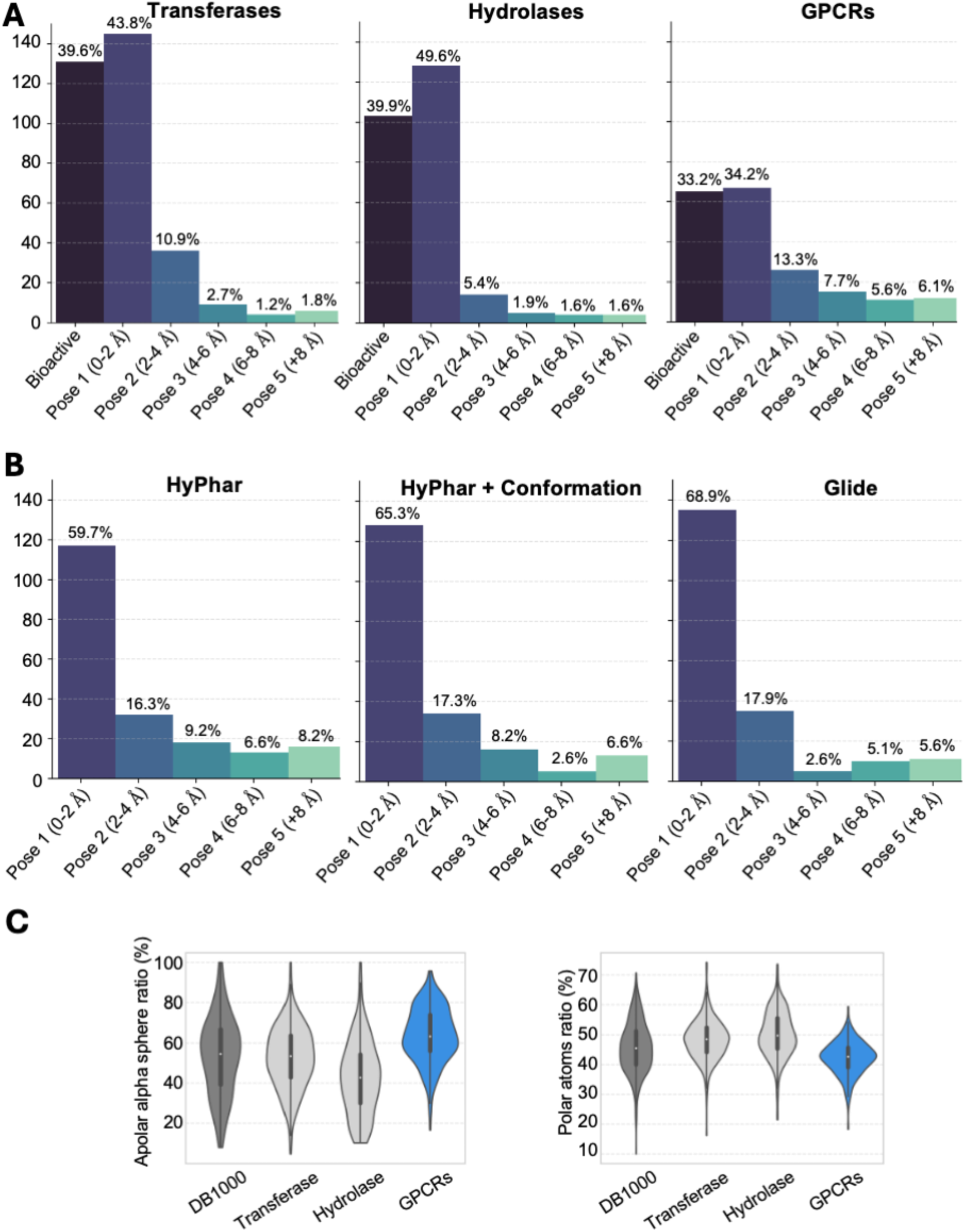
Effect of the nonpolar/polar balance in the binding pocket on the accuracy of *S*_*hydro*_. **(A)** Differences in accuracies for the 2 biggest families in the DB1000 dataset, compared to the DBGPCR. The accuracy of transferases (left), hydrolases (middle) and GPCRs are 83.4%, 89.5% and 67.4%, respectively. **(B)** Comparison of the accuracy of *S*_*hydro*_ and Glide (SP) for deviated poses only, excluding the bioactive pose. HyPhar accuracy on deviated poses for GPCRs set reaches a 59.7% of accuracy, which is increased up to 65,3% when adding the contribution of conformation term, in line with the increase in accuracy demonstrated in the manuscript. The combination of *S*_*hydro*_ and the conformational penalty reaches similar levels of accuracy as Glide scoring function (68.9%) despite the simplicity of the formalism used for both terms. **(C)** Differences in nonpolar alpha spheres and polar atoms between the DB1000 (grey), transferases (light grey), hydrolases (light grey) and GPCRs (blue). GPCRs show differences in the chemical descriptors of the pocket compared to the biggest DB1000 families.

The accuracy of *S*_*Hydro*_in discriminating the near-native pose for DBGPCR is slightly lower relative to the performance obtained for the 200 complexes included in the DB1000 test set (85.5%; see Fig. 3B). Indeed, comparison with the subset of complexes pertaining to transferases (331 complexes) and hydrolases (259 complexes) reveals a better performance for these two protein families (83.4% and 89.5%, respectively; Fig. 5B). However, this finding agrees with the preceding discussion about the dependence of the accuracy on the balance between nonpolar and polar patches of the binding cavity, as noted in the larger (lower) presence of nonpolar (polar) patches in the binding pockets of the DBGPCR compared with those of transferases and hydrolases complexes (Fig. 5C).

## Discussion

The results obtained for the distinct ligand-protein complexes examined above point out that the principle of hydropathic complementarity encoded in *S*_*hydro*_and *S*_*HINT*_is a promising strategy to discriminate between native-like and decoy poses of ligands. The two scoring functions benefit from the precise representation of both ligand and residues that shape the binding pocket, as well as from the usage of atomic descriptors that rely on a physically well-founded and consistent description of the hydropathic profile of the interacting partners. In the case of *S*_*hydro*_ this description is attained from first-principles QM calculations of the bioactive species of the ligand (and of the protein residues) using a finely parametrized version of the QM MST continuum solvation model. Regarding *S*_*HINT*_, the hydropathic description exploits atomic contributions to the LogP from experimentally measured values of small chemical moieties.

It should be noted that HINT was initially conceived by Abraham in the late 1980s, and implemented shortly thereafter^25,41^, assuming the hydropathic complementarity as a driving force of the recognition between ligand and protein, and its training data was, and remains, the relatively small set of LogP and fragment data compiled by Hansch and colleagues in the 1970s.^40^ While the applications of HINT progressed from protein crystallographic studies to rescoring of docking poses, and tools for hydropathic protein optimization and most recently protein structure predictions,^45,46^ the fundamentals and formalism have remained virtually unchanged for more than three decades. Thus, it is remarkable that the performance of *S*_*HINT*_ is close to that found for *S*_*hydro*_. This suggest that an updated training that exploits a more comprehensive set of experimental LogP data, supplemented with a tuning of the weights for the pairwise interactions included in HINT, could improve the predictive ability of *S*_*HINT*_.

The results also provide a glimpse of some methodological nuances that may enhance the predictive ability of the hydropathic complementarity scoring functions. First, the usage of conformation-weighted atomic descriptors for the protein residues can be improved by considering explicitly the conformational dependence of the hydropathicity of amino acids. Our previous studies^39^ showed that this is not relevant for nonpolar and aromatic residues, but the atomic contributions to the hydropathicity are more sensitive to the side chain conformation for polar and charged residues. On the other hand, given the uncertainties of force fields to estimate the conformational stress of small molecules, the implementation of more robust computational methods to predict the conformational penalty between poses may increase the weight of this energy component to ameliorate the discrimination between ligand’s poses. From a computational point of view, the cost of performing a QM continuum solvation calculation in water and *n*-octanol for the bioactive species of the ligand may limit the application of the hydropathic scoring to large chemical libraries this limitation can be largely alleviated via the introduction of artificial intelligence-models suitably trained to reproduce the conformation-dependent 3D hydropathy of drug-like compounds. This work, which is currently ongoing in our laboratory, will eventually enable a fast screening of chemical libraries.

Drug discovery is extremely complex; discriminating against the native-like pose of a drug-like candidate is just one of the many challenges that need to be faced in the race towards lead optimization. In this regard, the availability of the DB1000 and DBGPCR databases could be used by researchers as reference benchmark to assess novel developments in scoring functions. On the other hand, complementing the widely adopted description based on the enthalpic analysis of specific intermolecular interactions, exploiting the hydropathic complementarity between ligand and binding pocket provides an alternative way to disclose the key determinants that must be fulfilled by active compounds. In turn, since the hydropathic scoring function captures free energy components of ligand binding, it may be a solid foundation for developing structure-based predictors of binding affinity. Finally, two scenarios that deserve interest for future studies involve the extension to the presence of water-mediated contacts between ligand and receptor, which have not been explicitly considered for the complexes included in the DB1000 and DBGPCR datasets, and the effect of mutations at specific sites in the target protein, which may be relevant for exploring drug resistance mechanisms.

## Materials and Methods

### Datasets for benchmarking ligand-protein complementarity

#### Generation of ligand pose

The DB1000 dataset was prepared taking advantage of the structural information available in the PDBbind 2020 refined set, which encompass 5316 systems. DB1000 includes complexes with drug-like small molecules solved at a crystallographic resolution < 2.5 Å, and R-factor < 0.25, thus presumably lacking relevant structural issues. Duplicate structures for the same ligand-protein complex in the PDBbind 2020 were filtered out, retaining the entry with the highest resolution, and ligand-protein complexes where the ligand is coordinated to a metal center were excluded. The final selection of ligand-protein complexes was made to reflect the distribution of functional families according to their indexation in the Protein Data Bank (Fig. 1A).

Each of the 1000 systems included in the pose prediction dataset was prepared using the *PrepWizard* from *Schrödinger*, adding missing sidechains by using *Prime* and assigning the protonation state of the residues at pH 7.4 using *PROPKA*.^47–49^ The same pH was used to assign the protonation state to the ligands using *Epik*.^50^ After system preparation, Glide was used for the generation of the decoy poses of the ligand in Single Precision (SP) mode.^36^ At the end, besides the crystallographic pose, each system is linked to five ligand poses characterized by a positional RMSD relative to the heavy atoms of the X-ray pose in the ranges < 2, 2-4, 4-6, 6-8 and > 8Å (Fig. 2). The final pose prediction dataset consists of 1000 drug-like protein-ligand systems, containing a PDB file for the protein, an SDF file for the QM-parametrized X-ray poses and an additional SDF file for the five distinct poses.

#### Energy minimized dataset

With the aim of exploring the sensitivity of the results to geometrical refinement of the ligand pose, a refined subset of ligand-protein complexes was generated by minimizing the six ligand-protein complexes obtained for each protein using *Prime* with the OPLS4 force field without applying any restraints. Ligand-protein complexes that failed in retaining the proper distribution of ligand poses along the five distinct RMSD categories were excluded. This process led to a subset of 531 energy-minimized ligand-protein complexes, hereafter referred to as DBEMin, in which each protein is associated with the X-ray and five RMSD-ranked ligand poses.

#### G Protein-coupled receptor dataset

Keeping in mind the pharmacological relevance of GPCRs, the pose prediction protocol outlined above was used for generating ligand poses and evaluating the performance of the scoring function in this challenging class of receptors. To this end, a set of 196 ligand-GPCR complexes were extracted from the GPCRdb, ^51,52^ which is a comprehensive database with information of humanly curated structures. The choice of the selected structures was limited to cases where the ligand is bound to the orthosteric binding site of the GPCR. Furthermore, entries containing duplicated ligand-GPCR complexes were excluded, and GPCR complexes where a proper distribution of RMSD-ranked ligand poses could not be generated were eliminated. Overall, this procedure led to the generation of a dataset (DBGPCR) comprising 196 ligand-GPCR complexes.

### Hydropathic description of ligand-protein complexes

#### 3D atomic hydropathic (HyPhar) descriptors of ligands

Hyphar descriptors offer a three-dimensional view of molecular lipophilicity based on solvation theory. They are derived from QM calculations performed with the B3LYP/6-31G(d) parametrized version of the IEFPCM/MST continuum solvation model in water and *n*-octanol. The solvation free energies determined from these calculations are then combined to determine the *n*-octanol/water partition coefficient (logP). Remarkably, this approach allows performing QM calculations for the bioactive species, considering the specific chemical features (protonation, tautomerism, conformation, chirality) of the ligand in the bound state.

For our purposes here, it must be noted that the IEFPCM/MST can partition the solvation free energy, and hence the logP, into atomic contributions through application of a perturbative scheme of the coupling between the solute charge distribution and the solvent response.^53,54^ For a molecule (M) containing N atoms, this is achieved by decomposing the logP (or the transfer free energy, 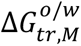) into electrostatic (*logP*_*ele*,*i*_), cavitation (*logP*_*cav*,*i*_) and van der Waals (*logP*_*VdW,i*_) components, which can be derived from the polar 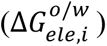 and nonpolar (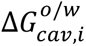 and *logP*_*VdW,i*_) contributions to the solvation free energy (Eqs. 3 and 4).

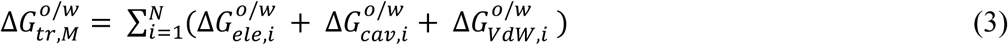

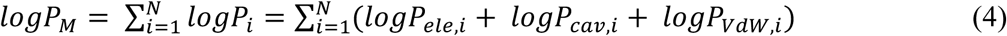

The electrostatic component can be partitioned into atomic contributions by applying a perturbative approximation that models the interaction between the solute’s charge distribution and the solvent’s reaction field (Eq. 5),

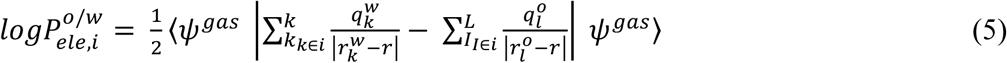

where *k* and *L* are the total number of reaction field charges in water 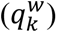 and *n*-octanol 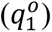, located in the positions 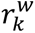 and 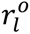, respectively, and *ψ*^g*as*^ denotes the wavefunction of the solute in the gas phase.

The atomic partitioning of the cavitation and van der Waals terms use their linear dependence to the solvent-accessible surface area of each atom in the molecule (Eqs. 6 and 7).

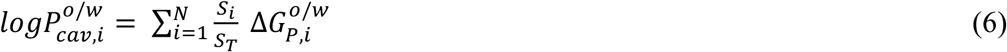

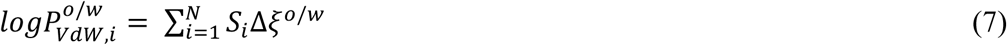

where 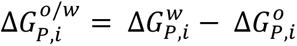, with Δ*G*_*P*,*i*_ is the cavitation free energy of the atom *i* and Δ*ξ*^*o*/w^ = *ξ*^w^ − *ξ*^*o*^, with *ξ* being the atomic surface tension, and *S*_*i*_ represents the atomic contribution to the total surface (*S*_*T*_).

This partitioning scheme provides a rigorous formalism for decomposition of the molecular logP into a 3D atomic map of hydrophobicity/hydrophilicity by assigning a contribution to each atom of the ligand.

Derivation of the ligand’s HyPhar descriptors were made via IEFPCM/MST B3LYP/6-31G(d) geometry optimizations of the X-ray pose in *n*-octanol, imposing restraints on the active torsions that define the ligand-bound conformation in the complex. This is convenient considering the sensitivity of the electron density (and hence the solvent response) to seemingly minor geometrical changes, especially in bond lengths and angles. Then, single-point calculations in the two solvents were performed to extract the hydrophobic parameters using the decomposition scheme into atomic contributions described above.

#### 3D atomic hydropathic (HyPhar) descriptors of protein

The description of the HyPhar descriptors for the protein residues exploits the protein-like hydrophobicity/hydrophilicity scale reported in a previous study. ^39^ This scale was conceived to provide pH-adjusted atomic contributions to the twenty natural amino acids from QM IEFPCM/MST calculations performed for the set of backbone-dependent conformational preferences of the residues. Briefly, this method considers both neutral and ionized forms of amino acids and incorporates p*K_a_* values to produce relevant 3D atomic distribution maps of the logP.

### The hydropathic scoring function

#### The hydropathic complementarity between ligand and protein

Eq. 1 provides a simple formalism to evaluate the hydrophobic/hydrophilic complementarity between ligand and protein. In this expression the atomic contributions of ligand and protein residues are determined from QM continuum solvation calculations that yield the ligand-based HyPhar descriptors (*logD*_*i*_), whereas the pH-dependent protein-like lipophilicity scale is adopted for the residues (*logD*_*j*_) in the target protein. The exponential function *e*^-*γiRi*j^ accounts for the distance-dependent decay between atoms *i* and *j*, which are separated at a distance *R*_*i*j_, and *γ*_*i*_ modulates the decay of the atomic pairwise interaction with the ligand’s atom.

The decay parameter *γ*_*i*_exhibits a sigmoidal dependence on the *logD*_*i*_value of atom *i* in the ligand. This balances interactions in which *logD*_*i*_ is very large, as may occur in the atomic hydrophobicity/hydrophilicity of charged atoms, which would bias the scoring towards certain types of pairwise contacts (i.e., salt bridges), whereas interactions for similar uncharged atoms would then have a negligible contribution to the total score. Moreover, *γ*_*i*_is also slightly adjusted to consider the distinct dependence of the geometrical distance of interacting atoms depending on the nature of the interaction with which the ligand’s atom is involved. Additionally, there is a scaling factor for *γ*_*i*_that adjusts its weight as a function of the ligand atomic logP value. This is done to reduce the contribution of interaction pairs where the ligand logP is very highly negative (e.g., charged atoms). The scaling is defined as a sigmoidal function (Eq. 8):

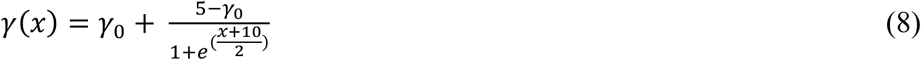

where *x* is the atomic ligand logP value and *γ*_M_ is the base scaling constant (Fig. 6).

**Figure 6.**
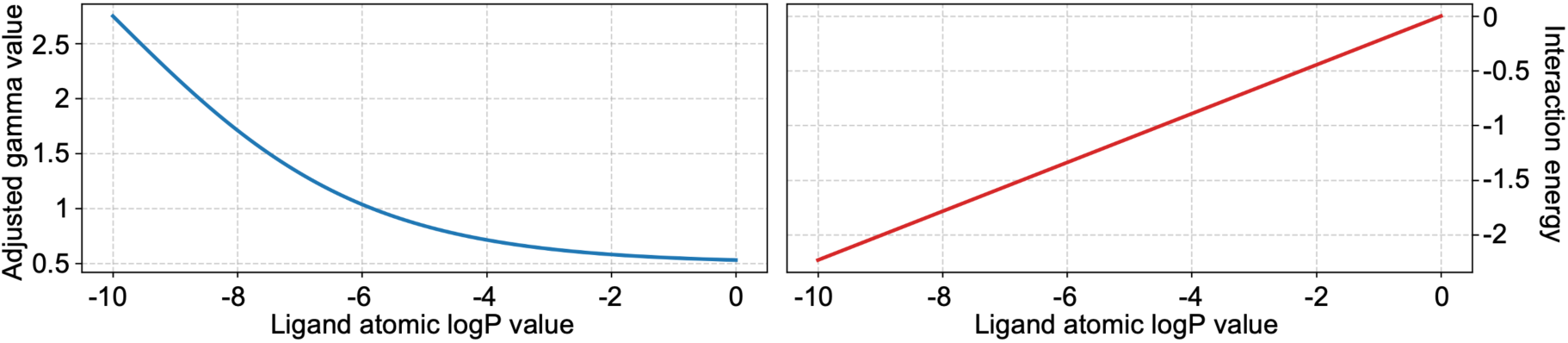
Scaling factor evolution and decay of logP values. (Left) Adjustment of the decay parameter γ(x) as a function of the ligand atomic logP value, shown for a baseline γ_0_=0.5. (Right) Corresponding interaction energy as a function of ligand atomic logP (assuming a protein logP value of 1).

### Refinements to the scoring function

The effect of supplementing the hydrophobic/hydrophilic scoring function (*S*_*hydro*_) with other physical contributions on the prediction of the bioactive pose was exploring considering four factors, which include the conformational stress of the ligand, the total desolvation free energy upon binding of the ligand, and the categorization of the distinct interactions that mediate the binding between ligand and protein (Eq. 9).

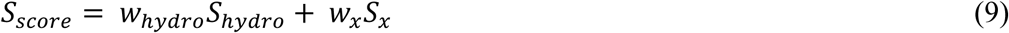

where x stands for the contributions due to conformational energy (*conf*), desolvation (*solv*) and number of pairwise interactions (*int*) of the distinct ligand poses, *S*_*conf*_ is the conformational energy score of the ligand, *S*_*solv*_ is the solvation effect score of the protein-ligand complex, and *S*_*int*_ is the sum of protein-ligand interactions score, and *w*_*hydro*_, *w*_*conf*_, *w*_*solv*_, and *w*_*int*_ represent the weights of each term to the final *S*_*score*_.

#### Conformational term

The *S*_*conf*_ term was computed using the MMFF94s force field as implemented in the RDKit toolkit. ^55^ The conformational penalty was estimated from the difference between the energy of the partially optimized ligand pose (keeping active torsions frozen at the crystallographic values) and the closest conformational minimum obtained from full optimization of the ligand. Even though this is a crude approximation, which otherwise would require a complete conformational sampling to find the global minima, this scheme was already used in previous studies to estimate the conformational penalty.

#### Solvation term

Binding of the ligand may lead to a complete burial into the pocket or a partial occlusion, with part of the ligand being exposed to the bulk solvent in this latter case. Similarly, water molecules filling the binding pocket may be fully displaced upon ligand binding or some subpockets may still retain the interactions with water molecules. Accordingly, this term was introduced to examine the potential contribution of the desolvation of the ligand and protein using Eq. 10.

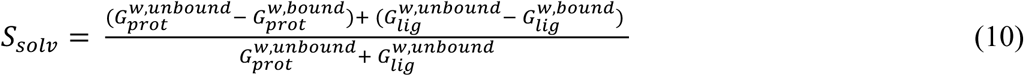

where 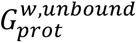 and 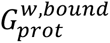 represent the desolvation free energy of the protein in the unbound and bound states, respectively, and 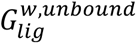 and 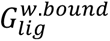 stands for corresponding contributions due to the ligand (Eqs. 11-14).

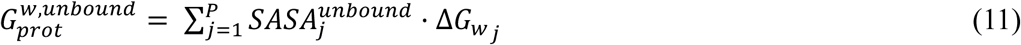

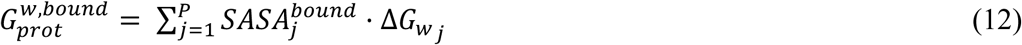

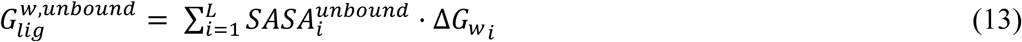

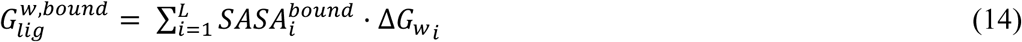

*w*here 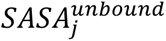 and 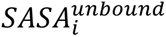 represent the solvent accessible surface area j(SASA) of atom *j* (protein) or *i* (ligand), respectively, in the unbound state, and 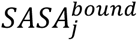 and 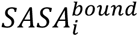 denote the SASA of protein and ligand, respectively, in the bound state.

#### Interaction term

Finally, we have explored the addition of a correction term that collects a qualitative description of the interactions formed between the ligand and the protein, as shown in Eq. 15.

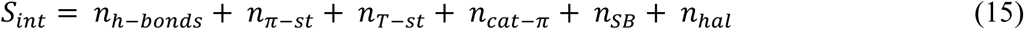

where the terms in Eq. 15 reflects the number of hydrogen bonds (*n*_*h*–*bonds*_ ), aromatic *π*-*π* and T stacking (*n*_*π*–*st*_, *n*_*T*–*st*_), cation-*π* contacts (*n*_*cat*–*π*_), salt bridge (*n*_*SB*_), and halogen-bond interactions (*n*_*hal*_).

To this end, the introduction of corrections for the formation of these interactions takes place subject to the fulfilment of the geometric parameters and cutoffs defined for each kind of interaction between ligand and protein (Fig. 7).

**Figure 7.**
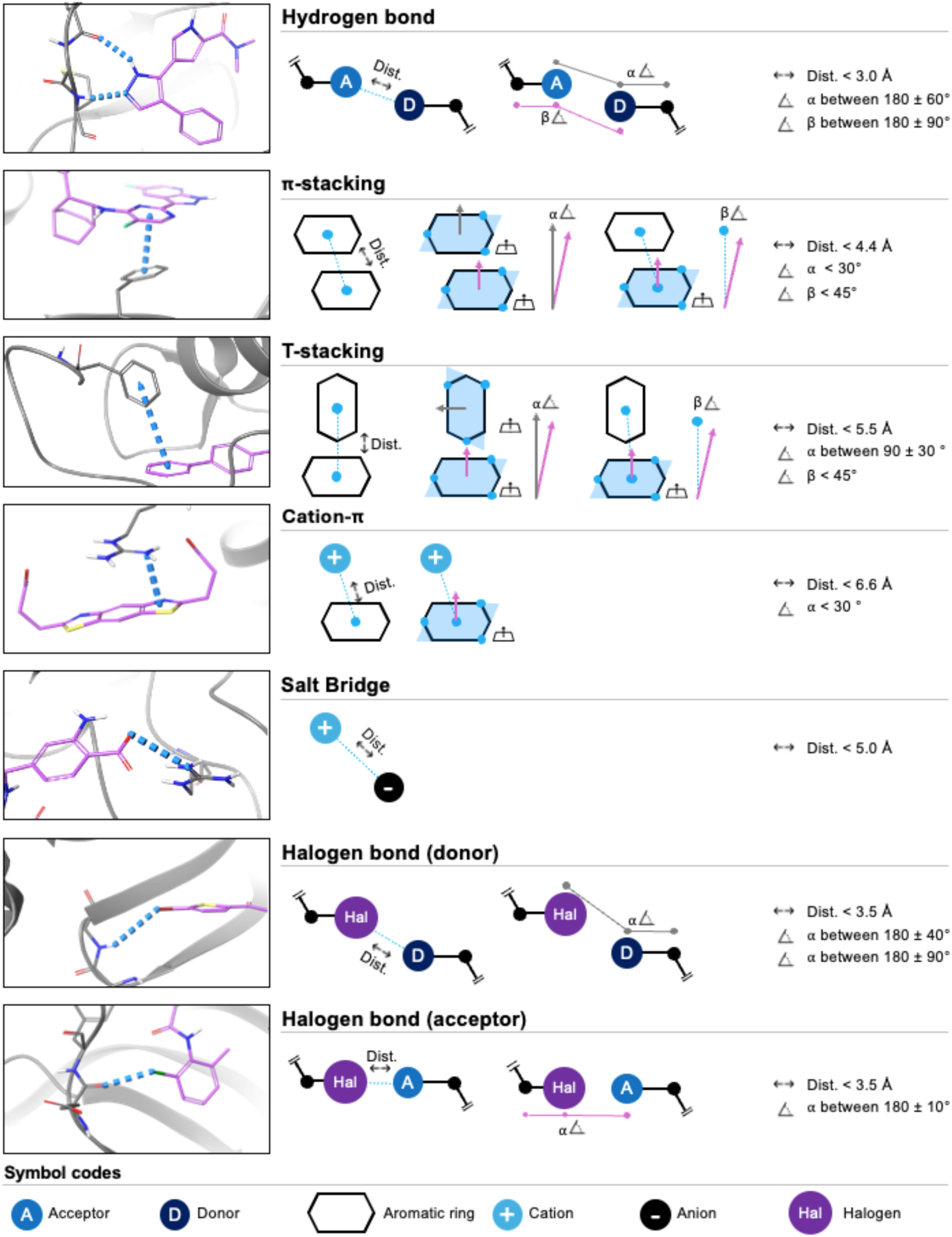
Schema of the interaction cutoffs to define the interaction term. The interaction term consists of the sum of the total number of interactions between the ligand and the protein.

#### Weights of the scoring function

Optimization of the weights for each component in Eq. 9 was performed by normalizing the values of *S*_*hydro*_, *S*_*conf*_, *S*_*solv*_, and *S*_*int*_, so that each variable was adjusted to have a mean of zero and a standard deviation of one unit. The weights *w*_*hydro*_, *w*_*conf*_ and *w*_*solv*_ were then determined using a Bayesian optimization with the goal of maximizing the accuracy of the final model in finding the native pose.

To this end, we randomly split our dataset of ligand–protein complexes into a training set (800 systems) and a test set (200 systems). The objective function was described as the accuracy in finding the bioactive or the lowest deviated pose (0-2 Å) among the best scores for all the system in the training set. We used the Scikit-Optimize package, and we set up the optimizer to explore the parameter space over 100 calls per iteration, with 25 random initializations and 5 restarts of the local optimizer.^56^ Convergence was evaluated using an early stopping rule that considered slope and variance from recent iterations. Results from all the runs are provided in Supplementary Material Fig. S5.

To check if the training and test split was appropriate, we conducted a 5-fold cross-validation on the pose prediction dataset and computed the docking accuracies for the hydrophobic term on each test set (200 systems). The docking accuracies were 83, 85, 82, and 83.5, with an average of 84.0 ± 1.7. We also adjusted the scoring function weights for each fold to see if they were like the selected test. The results shown that the weights obtained for each pair of variables obtained by combining *S*_*hydro*_ separately with the other terms (*S*_*conf*_, *S*_*solv*_, and *S*_*int*_) were very similar (Supplementary Material Fig. S7).

#### Evaluation metrics and optimizations

To estimate the performance of the scoring function in discriminating native (or near native) poses from highly deviated ones, the docking accuracy was estimated as the percentage of systems in which the X-ray pose or the pose with the near-native pose (RMSD < 2Å) is assigned the best scoring value. This metric provides a quantitative measure of how precisely the scoring function can identify the most accurate binding pose, and hence its ability to prioritize near-native poses over those that are significantly deviated from the experimentally determined structures. High docking accuracy indicates that the scoring function is reliable in recognizing the correct pose.

## Data availability

All the datasets (DB1000, DBEMin and DBGPCR), including the X-ray and decoy poses and hydropathic descriptors of ligands, are publicly available in Zenodo (https://zenodo.org/records/17294819).

## Code availability

The source code used in this study will be made available upon request to the corresponding authors.

## Author contributions

B.M.-L. designed the framework and implemented the code. W.J.Z. provided support in implementing the protein Hyphar parameters in the code. E.H., G.E.K., J.S., J.V. and F.J.L. contributed conceptually to the method development and provided supervision throughout the project. All authors edited and reviewed the manuscript.

## Acknowledgments

B.M.-L. acknowledges the support from the Generalitat de Catalunya through the Industrial Doctorate PhD fellowship (AGAUR 2022DI112). This study was funded by the Spanish Ministerio de Ciencia e Innovación (PID2020-117646RB-I00 MCIN/AEI/10.13039/501100011033; Maria de Maetzu CEX2021-001202-M), the Generalitat de Catalunya (2021SGR00671), and the Consorci de Serveis Universitaris de Catalunya (CSUC) is acknowledged for computational facilities (Molecular Recognition project).

## Competing interests

B.M.-L., E.H. and J.V. work in Pharmacelera SL and F.J.L. is scientific advisor of this company but they all declare no financial competing interests. All other authors declare no financial or non-financial competing interests.

---

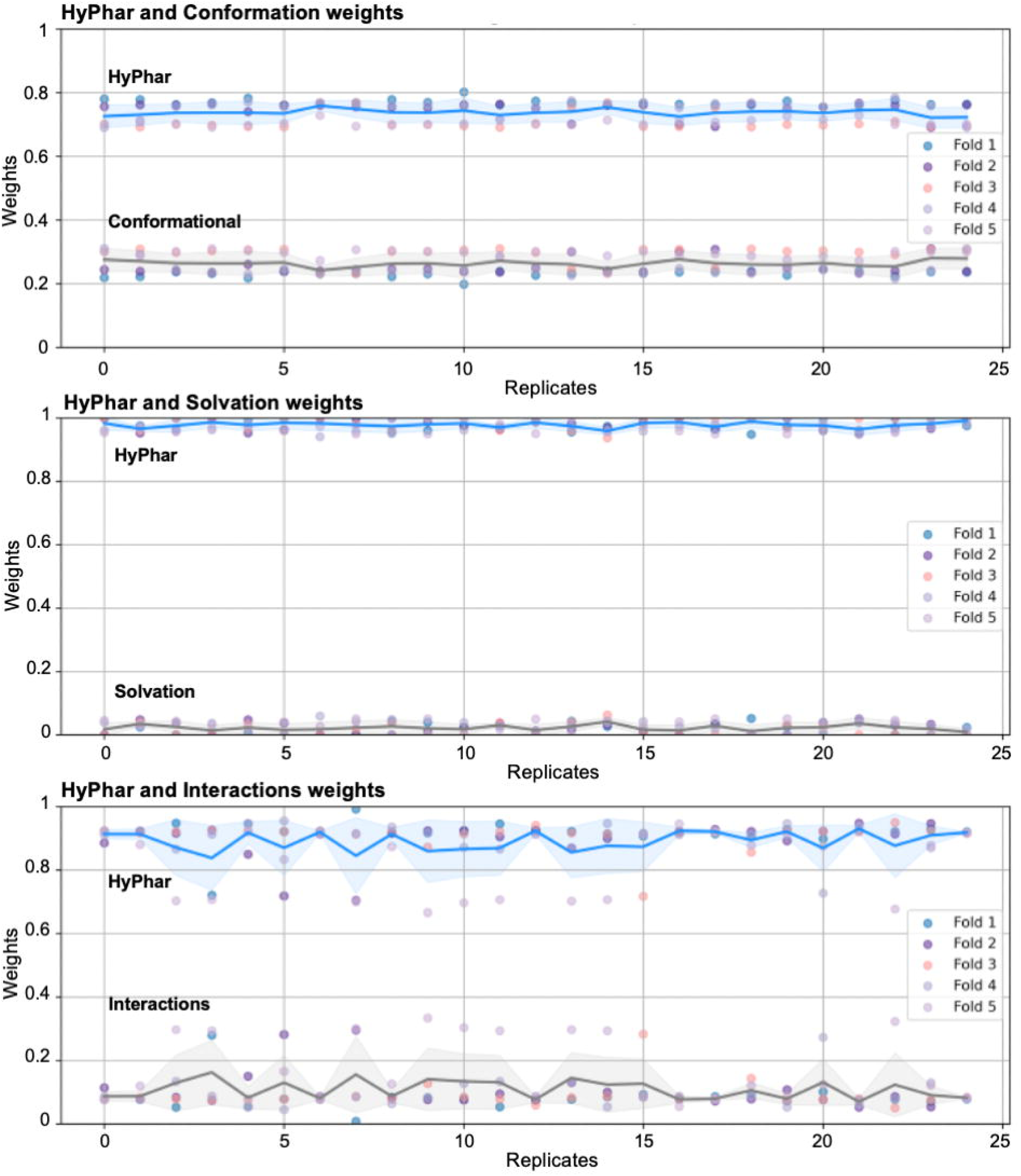

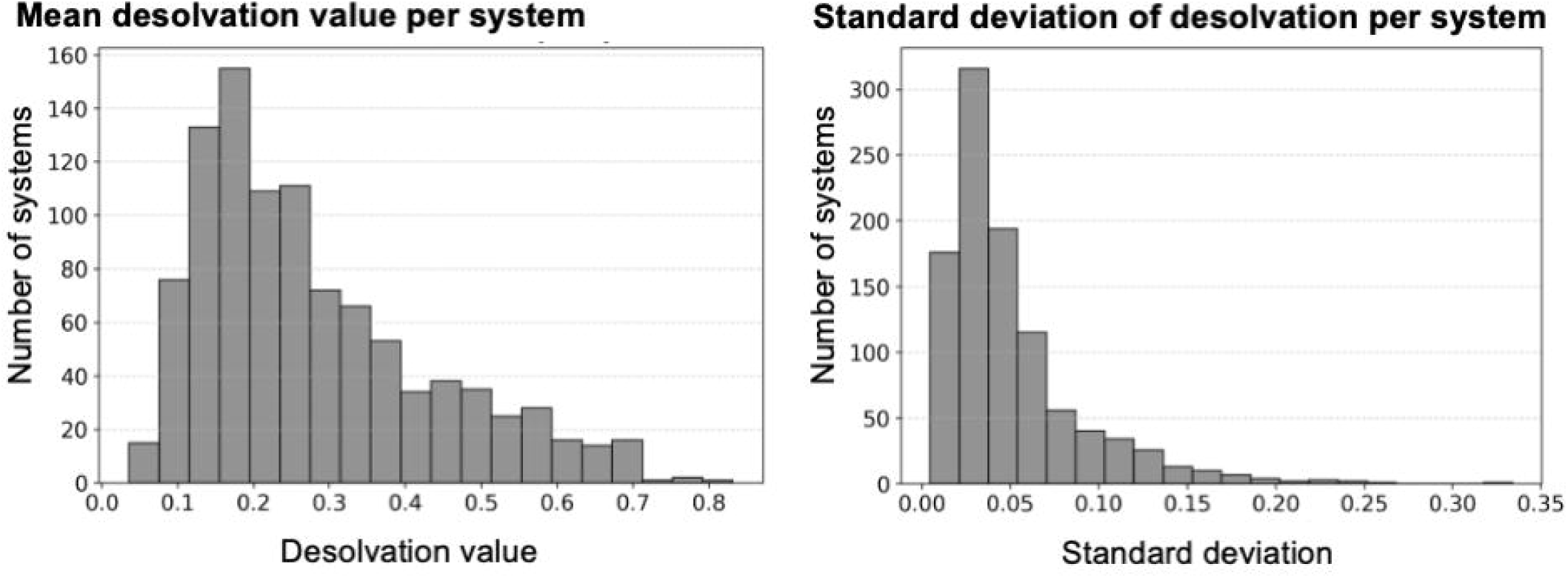

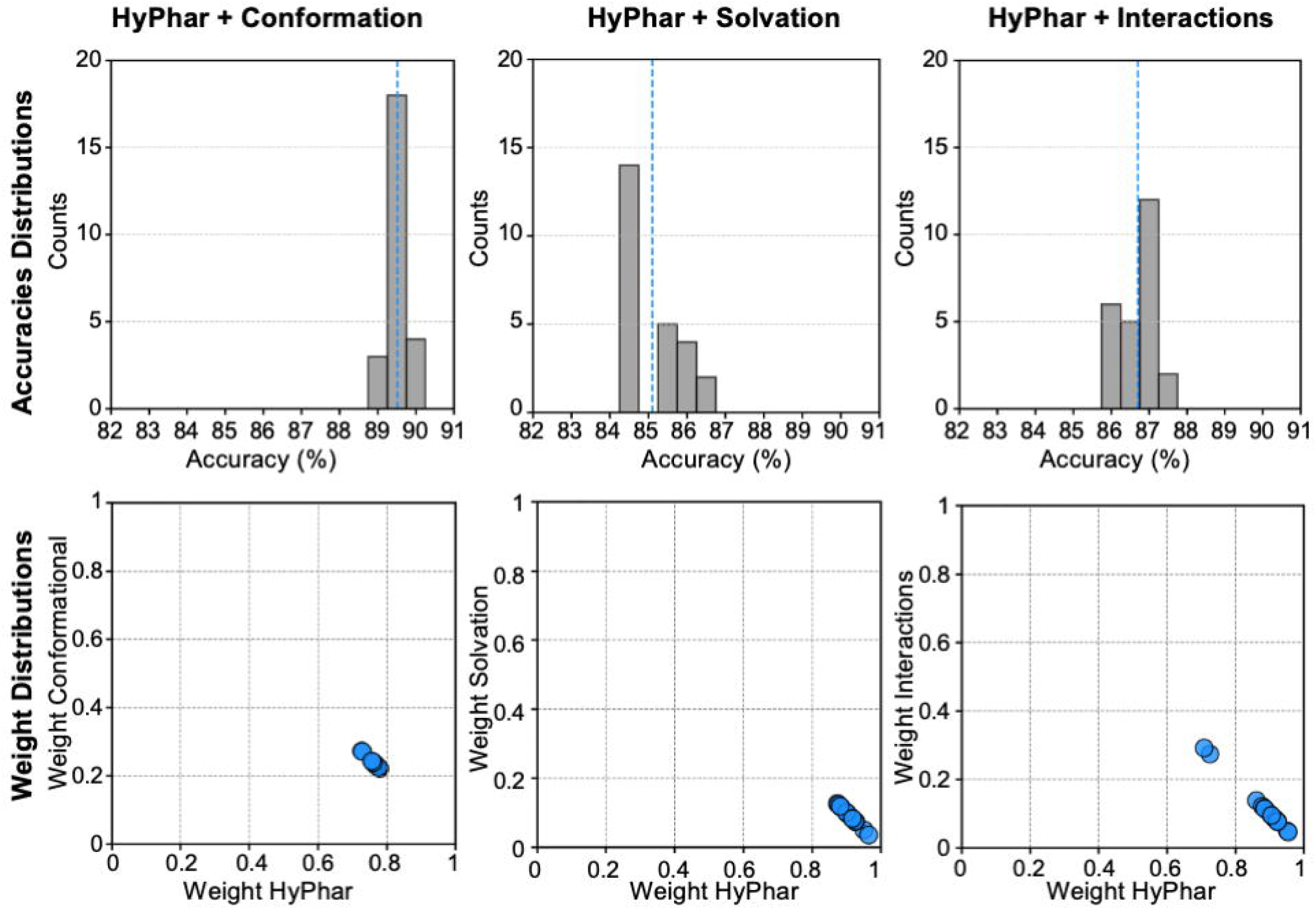

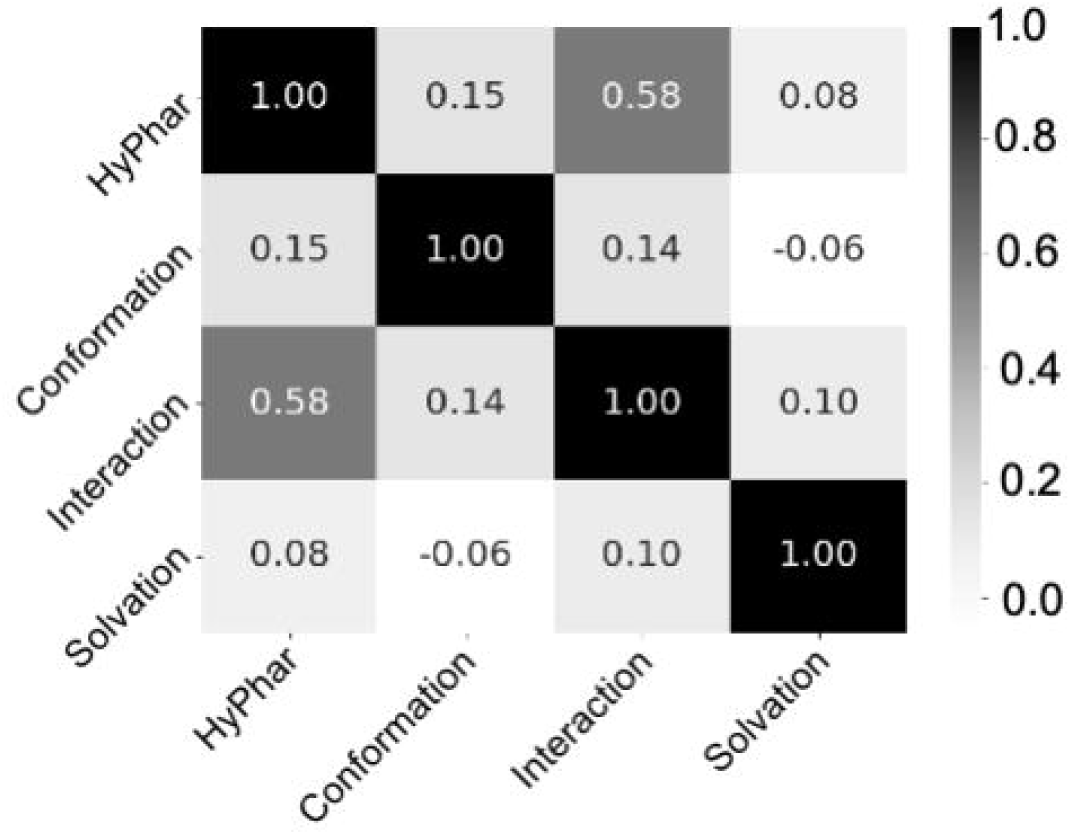

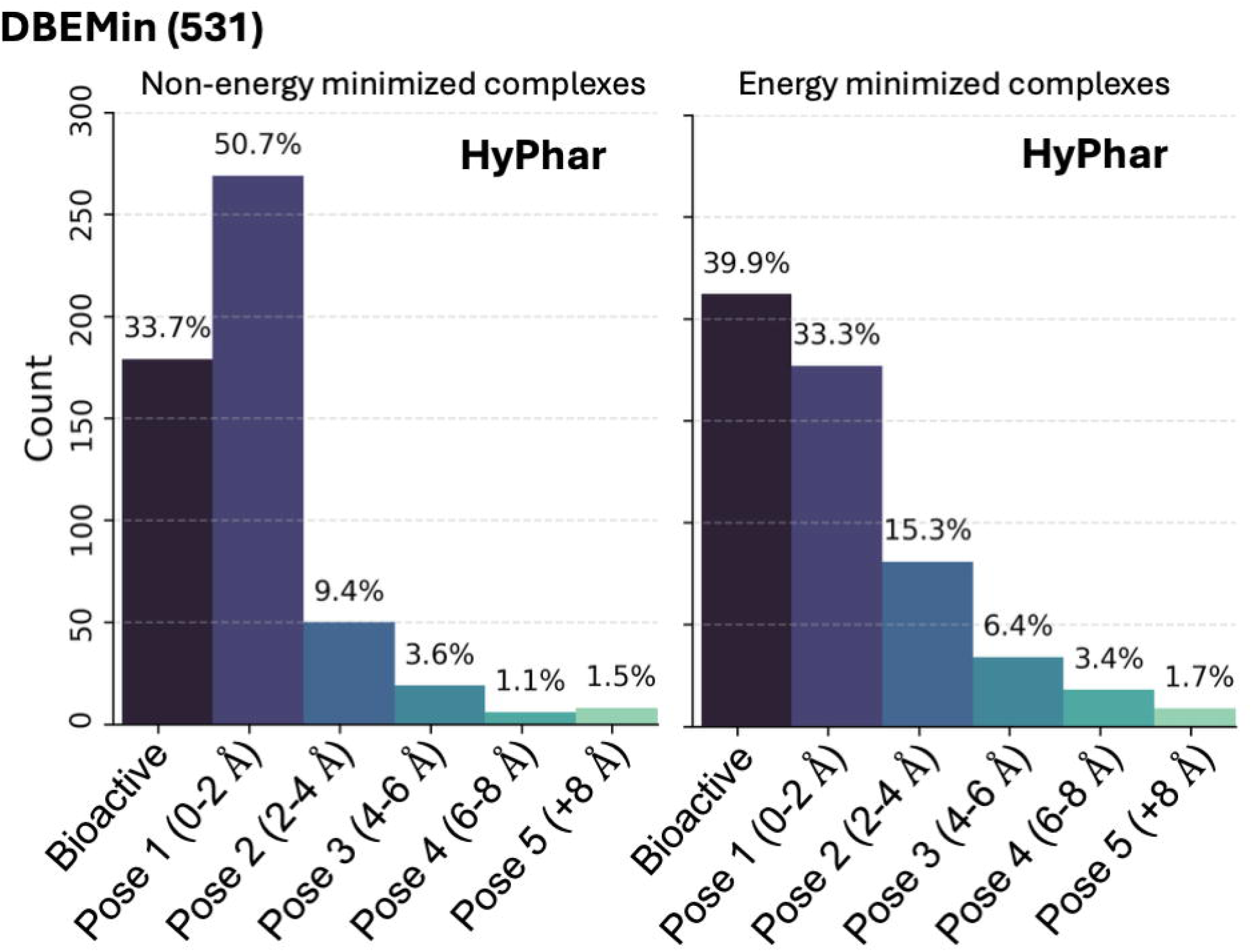

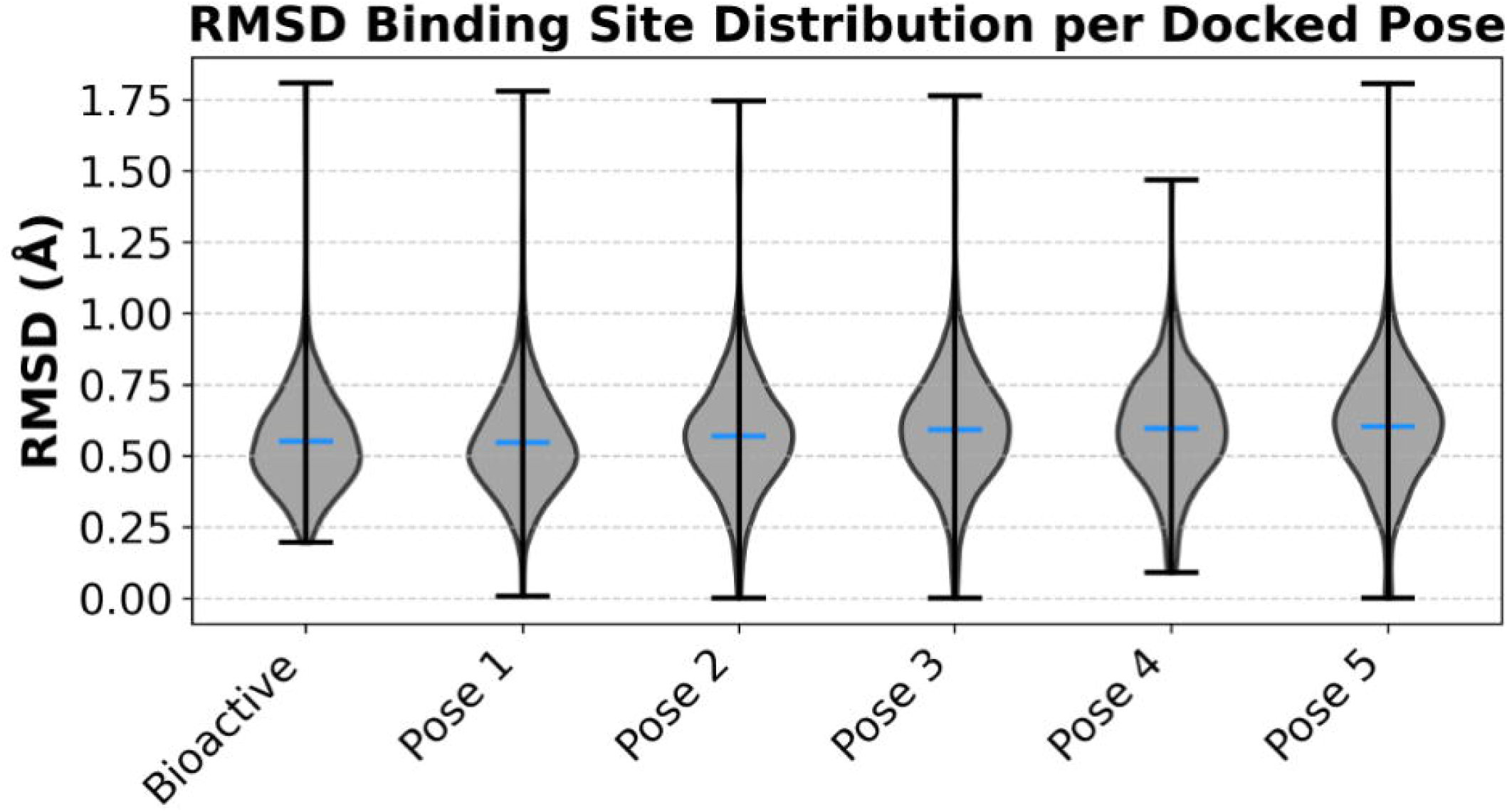

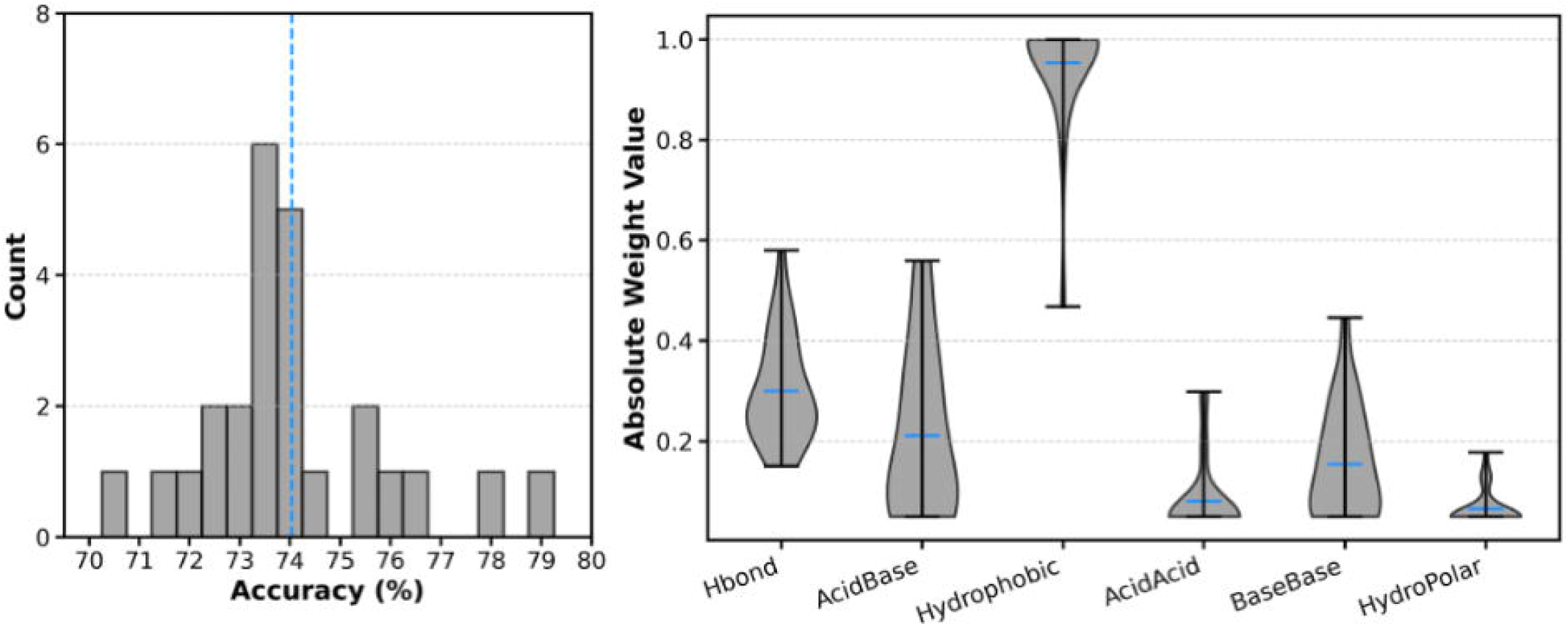

## Notes

https://zenodo.org/records/17294819

### Competing Interest Statement

The authors have declared no competing interest.

## Bibliography

1. Bissantz, C., Kuhn, B. & Stahl, M. A medicinal chemist’s guide to molecular interactions. J Med Chem 53, 5061–5084 (2010).

2. Approaches to the description and prediction of the binding affinity of small-molecule ligands to macromolecular receptors - PubMed. https://pubmed.ncbi.nlm.nih.gov/12203463/.

3. Ferenczy, G. G. & Keseru, G. M. Thermodynamics guided lead discovery and optimization. Drug Discov Today 15, 919–932 (2010).

4. Reymond, J. L. The Chemical Space Project. Acc Chem Res 48, 722–730 (2015).

5. Lyu, J. et al. Ultra-large library docking for discovering new chemotypes. Nature 566, 224–229 (2019).

6. Kuan, J., Radaeva, M., Avenido, A., Cherkasov, A. & Gentile, F. Keeping pace with the explosive growth of chemical libraries with structure-based virtual screening. Wiley Interdiscip Rev Comput Mol Sci 13, e1678 (2023).

7. Cherkasov, A. The ‘Big Bang’ of the chemical universe. Nat Chem Biol 19, 667–668 (2023).

8. Sindt, F., Bret, G. & Rognan, D. On the Difficulty to Rescore Hits from Ultralarge Docking Screens. J Chem Inf Model 65, 5553–5566 (2025).

9. Luo, J., Wei, W., Waldispühl, J. & Moitessier, N. Challenges and current status of computational methods for docking small molecules to nucleic acids. Eur J Med Chem 168, 414–425 (2019).

10. Cavasotto, C. & W. Orry, A. Ligand docking and structure-based virtual screening in drug discovery. Curr Top Med Chem 7, 1006–1014 (2007).

11. Paggi, J. M., Pandit, A. & Dror, R. O. The Art and Science of Molecular Docking. Annu Rev Biochem 93, 389–410 (2024).

12. Guedes, I. A., Pereira, F. S. S. & Dardenne, L. E. Empirical scoring functions for structure-based virtual screening: Applications, critical aspects, and challenges. Front Pharmacol **9**, 411637 (2018).

13. Vittorio, S. et al. Addressing docking pose selection with structure-based deep learning: Recent advances, challenges and opportunities. Comput Struct Biotechnol J 23, 2141–2151 (2024).

14. Yim, J. et al. Diffusion models in protein structure and docking. Wiley Interdiscip Rev Comput Mol Sci 14, e1711 (2024).

15. Cavasotto, C. N. & Aucar, M. G. High-Throughput Docking Using Quantum Mechanical Scoring. Front Chem 8, 531964 (2020).

16. Majewski, M., Ruiz-Carmona, S. & Barril, X. Dynamic Undocking: A Novel Method for Structure-Based Drug Discovery. Methods Mol Biol 1824, 195–216 (2018).

17. Guo, Z. et al. Identification of Protein–Ligand Binding Sites by the Level-Set Variational Implicit-Solvent Approach. J Chem Theory Comput 11, 753–765 (2015).

18. Sánchez-Cruz, N., Medina-Franco, J. L., Mestres, J. & Barril, X. Extended connectivity interaction features: improving binding affinity prediction through chemical description. Bioinformatics 37, 1376–1382 (2021).

19. Fauman, E. B., Rai, B. K. & Huang, E. S. Structure-based druggability assessment--identifying suitable targets for small molecule therapeutics. Curr Opin Chem Biol 15, 463–468 (2011).

20. Schmidtke, P. & Barril, X. Understanding and predicting druggability. A high-throughput method for detection of drug binding sites. J Med Chem 53, 5858–5867 (2010).

21. Hajduk, P. J., Huth, J. R. & Fesik, S. W. Druggability indices for protein targets derived from NMR-based screening data. J Med Chem 48, 2518–2525 (2005).

22. Cheng, A. C. et al. Structure-based maximal affinity model predicts small-molecule druggability. Nat Biotechnol 25, 71–75 (2007).

23. Young, T., Abel, R., Kim, B., Berne, B. J. & Friesner, R. A. Motifs for molecular recognition exploiting hydrophobic enclosure in protein–ligand binding. Proceedings of the National Academy of Sciences 104, 808–813 (2007).

24. Sheridan, R. P., Maiorov, V. N., Holloway, M. K., Cornell, W. D. & Gao, Y. D. Drug-like density: A method of quantifying the ‘bindability’ of a protein target based on a very large set of pockets and drug-like ligands from the protein data bank. J Chem Inf Model 50, 2029–2040 (2010).

25. Eugene Kellogg, G. & Abraham, D. J. Hydrophobicity: is LogPo/w more than the sum of its parts? Eur J Med Chem 35, 651–661 (2000).

26. Kellogg, G. E., Marabotti, A., Spyrakis, F. C Mozzarelli, A. HINT, a code for understanding the interaction between biomolecules: a tribute to Donald J. Abraham. Front Mol Biosci 10, 1194962 (2023).

27. Caron, G. et al. Steering New Drug Discovery Campaigns: Permeability, Solubility, and Physicochemical Properties in the bRo5 Chemical Space. ACS Med Chem Lett 12, 13 (2021).

28. Sternicki, L. M. & Poulsen, S. A. Fragment-based drug discovery campaigns guided by native mass spectrometry. RSC Med Chem 15, 2270 (2024).

29. Vázquez, J. et al. Screening and Biological Evaluation of Soluble Epoxide Hydrolase Inhibitors: Assessing the Role of Hydrophobicity in the Pharmacophore-Guided Search of Novel Hits. J Chem Inf Model 63, 3209–3225 (2023).

30. Nesbitt, N. M. et al. Small molecule BLVRB redox inhibitor promotes megakaryocytopoiesis and stress thrombopoiesis in vivo. Nature Communications 2025 1C:1 16, 1–18 (2025).

31. Luque, F. J. et al. Continuum solvation models: Dissecting the free energy of solvation. Physical Chemistry Chemical Physics 5, 3827–3836 (2003).

32. Vázquez, J. et al. Development and Validation of Molecular Overlays Derived from Three-Dimensional Hydrophobic Similarity with PharmScreen. J Chem Inf Model 58, 1596–1609 (2018).

33. Ginex, T. et al. Development and validation of hydrophobic molecular fields derived from the quantum mechanical IEF/PCM-MST solvation models in 3D-QSAR. J Comput Chem 37, 1147–1162 (2016).

34. Liu, Z. et al. Forging the Basis for Developing Protein–Ligand Interaction Scoring Functions. Acc Chem Res 50, 302–309 (2017).

35. Berman, H. M. et al. The Protein Data Bank. Nucleic Acids Res 28, 235–242 (2000).

36. Friesner, R. A. et al. Glide: A New Approach for Rapid, Accurate Docking and Scoring. 1. Method and Assessment of Docking Accuracy. J Med Chem 47, (2004).

37. Curutchet, C., Orozco, M. & Luque, F. J. Solvation in octanol: parametrization of the continuum MST model. J Comput Chem 22, 1180–1193 (2001).

38. Soteras, I., Curutchet, C., Bidon-Chanal, A., Orozco, M. & Javier Luque, F. Extension of the MST model to the IEF formalism: HF and B3LYP parametrizations. Journal of Molecular Structure: THEOCHEM 727, 29–40 (2005).

39. Zamora, W. J., Campanera, J. M. & Luque, F. J. Development of a structure-based, pH-Dependent lipophilicity scale of amino acids from continuum solvation calculations. Journal of Physical Chemistry Letters 10, 883–889 (2019).

40. Leo, A., Hansch, C. & Elkins, D. Partition coefficients and their uses. Chem Rev 71, 525–616 (2002).

41. Kellogg, G. E., Semus, S. F. & Abraham, D. J. HINT: A new method of empirical hydrophobic field calculation for CoMFA. J Comput Aided Mol Des 5, (1991).

42. Lu, C. et al. OPLS4: Improving Force Field Accuracy on Challenging Regimes of Chemical Space. J Chem Theory Comput 17, 4291–4300 (2021).

43. Merck molecular force field. I. Basis, form, scope, parameterization, and performance of MMFF94 - Halgren - 1996 - Journal of Computational Chemistry - Wiley Online Library. https://onlinelibrary.wiley.com/doi/10.1002/(SICI)1096-987X(199604)17:5/6%3C490::AID-JCC1%3E3.0.CO;2-P.

44. Le Guilloux, V., Schmidtke, P. C Tuffery, P. Fpocket: An open source platform for ligand pocket detection. BMC Bioinformatics 10, 1–11 (2009).

45. Al Mughram, M. H. et al. 3D Interaction Homology: Hydropathic Analyses of the “π–Cation” and “π–π” Interaction Motifs in Phenylalanine, Tyrosine, and Tryptophan Residues. J Chem Inf Model 61, 2937–2956 (2021).

46. Agosta, F., Kellogg, G. E. & Cozzini, P. From oncoproteins to spike proteins: the evaluation of intramolecular stability using hydropathic force field. J Comput Aided Mol Des 36, 797–804 (2022).

47. Protein Preparation Workflow - Schrödinger. https://www.schrodinger.com/life-science/learn/white-papers/protein-preparation-workflow/.

48. Prime | Schrödinger. https://www.schrodinger.com/platform/products/prime/.

49. Olsson, M. H. M., SØndergaard, C. R., Rostkowski, M. & Jensen, J. H. PROPKA3: Consistent Treatment of Internal and Surface Residues in Empirical pKa Predictions. J Chem Theory Comput 7, 525–537 (2011).

50. Johnston, R. C. et al. Epik: p Ka and Protonation State Prediction through Machine Learning. J Chem Theory Comput 19, 2380–2388 (2023).

51. Herrera, L. P. T. et al. GPCRdb in 2025: adding odorant receptors, data mapper, structure similarity search and models of physiological ligand complexes. Nucleic Acids Res 53, D425–D435 (2025).

52. Pándy-Szekeres, G. et al. GPCRdb in 2018: adding GPCR structure models and ligands. Nucleic Acids Res 46, D440–D446 (2018).

53. Javier Luque, F., Barril, X. & Orozco, M. Fractional description of free energies of solvation. J Comput Aided Mol Des 13, 139–152 (1999).

54. Luque, F. J., Bofill, J. M. & Orozco, M. New strategies to incorporate the solvent polarization in self-consistent reaction field and free-energy perturbation simulations. J Chem Phys 103, 10183–10191 (1995).

55. An overview of the RDKit — The RDKit 2025.03.6 documentation. https://www.rdkit.org/docs/Overview.html.

56. Pedregosa Fabianpedregosa, F. et al. Scikit-learn: Machine Learning in Python. The Journal of Machine Learning Research 12, 2825–2830 (2011).

