## Supplementary Material for "Assessing the ligand native-like pose using a quantum mechanical-derived hydropathic score for protein-ligand complementarity"

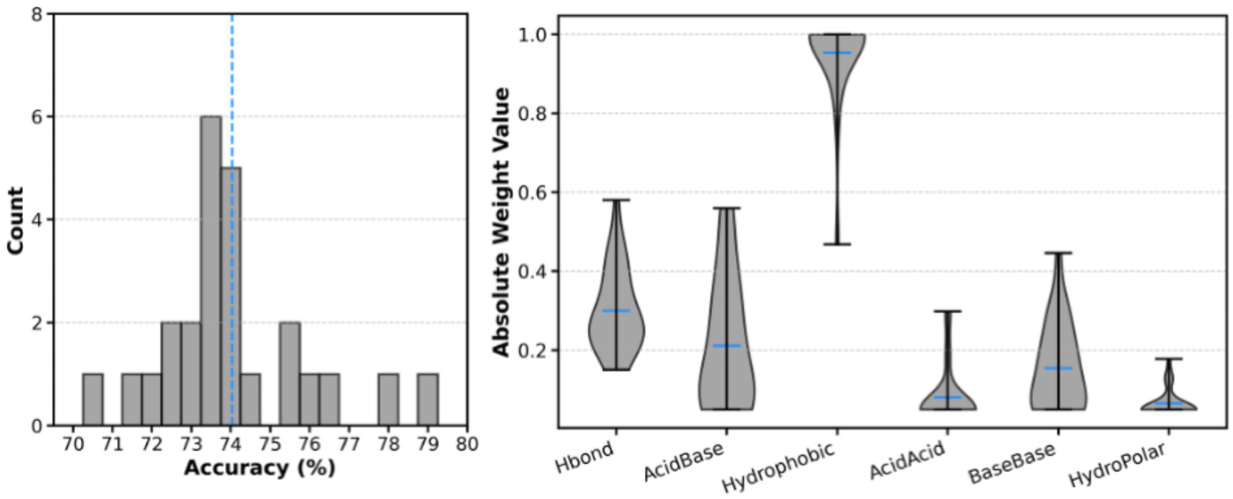


**Fig. S1. Results of the Bayesian optimization of the pairwise interaction components in *$S_{HINT}$*.** (Left) Distribution of the accuracy obtained in 25 Bayesian optimization calculations. The dashed blue line represents the accuracy averaged for the distinct calculations. (Right) Violin plots showing the distribution of the weights obtained for hydrophobic, hydrogen-bond, acid-base, base-base, acid-acid and hydrophobic-polar interactions in the distinct optimization calculations.

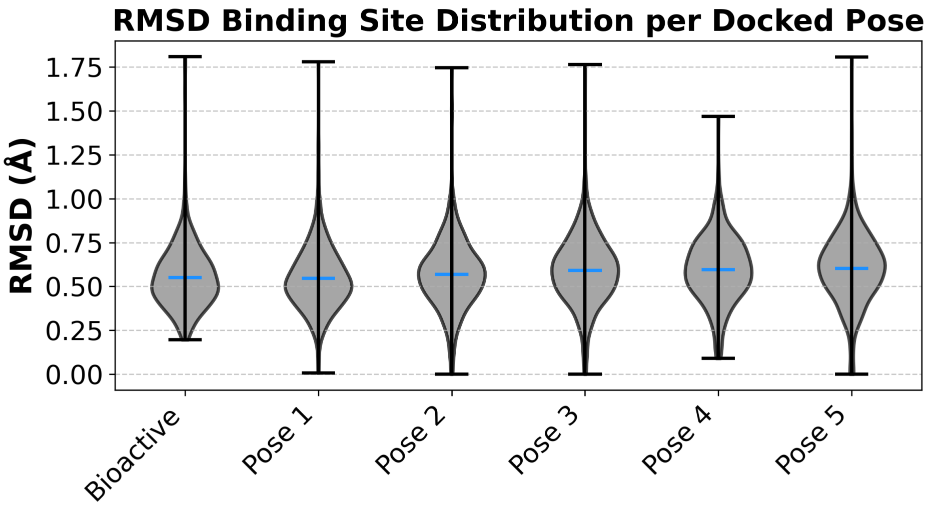


**Fig. S2. Geometrical changes introduced in the residues that shape the binding pocket upon energy minimization with the OPLS4 force field.** The violin plots show the distribution of positional root-mean square deviations of the heavy atoms for the residues in the binding pocket determined relative to their position in the X-ray crystallographic structure. The energy minimization was performed for the complexes formed with ligands in either the X-ray pose or the docked poses.

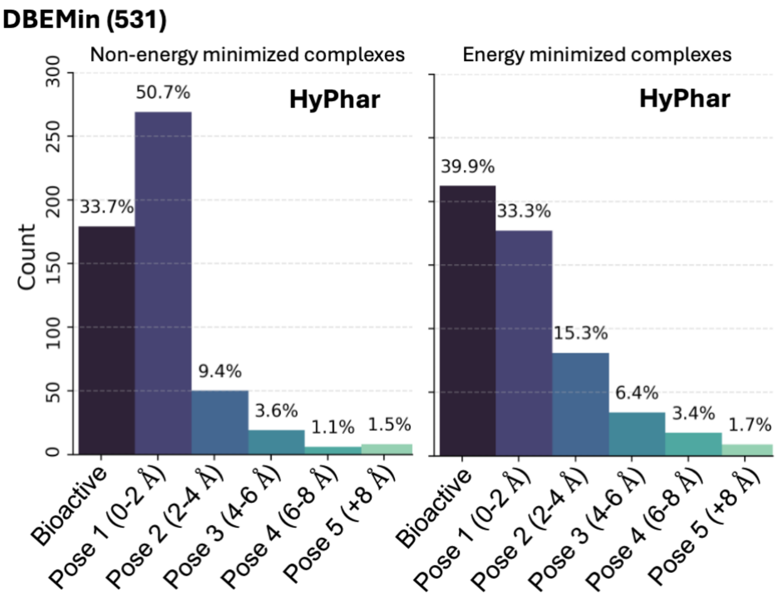


**Fig. S3. Effect of the geometrical refinement of the ligand-protein complexes on the performance of $S_{HYDRO}$ for 531 targets included in the DBEMin dataset**. Left and right plots denote the distribution of poses obtained with and without energy-minimization.

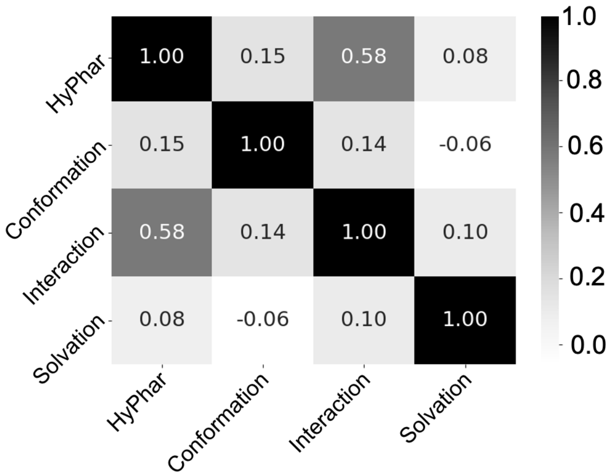


**Fig. S4. Correlation between selected properties determined for the distinct poses of the ligand in the set of ligand-protein complexes.** Properties include the hydrophobicity score, the conformational stress, the desolvation penalty, and the number of specific interactions formed between the ligand and the residues in the protein cavity.

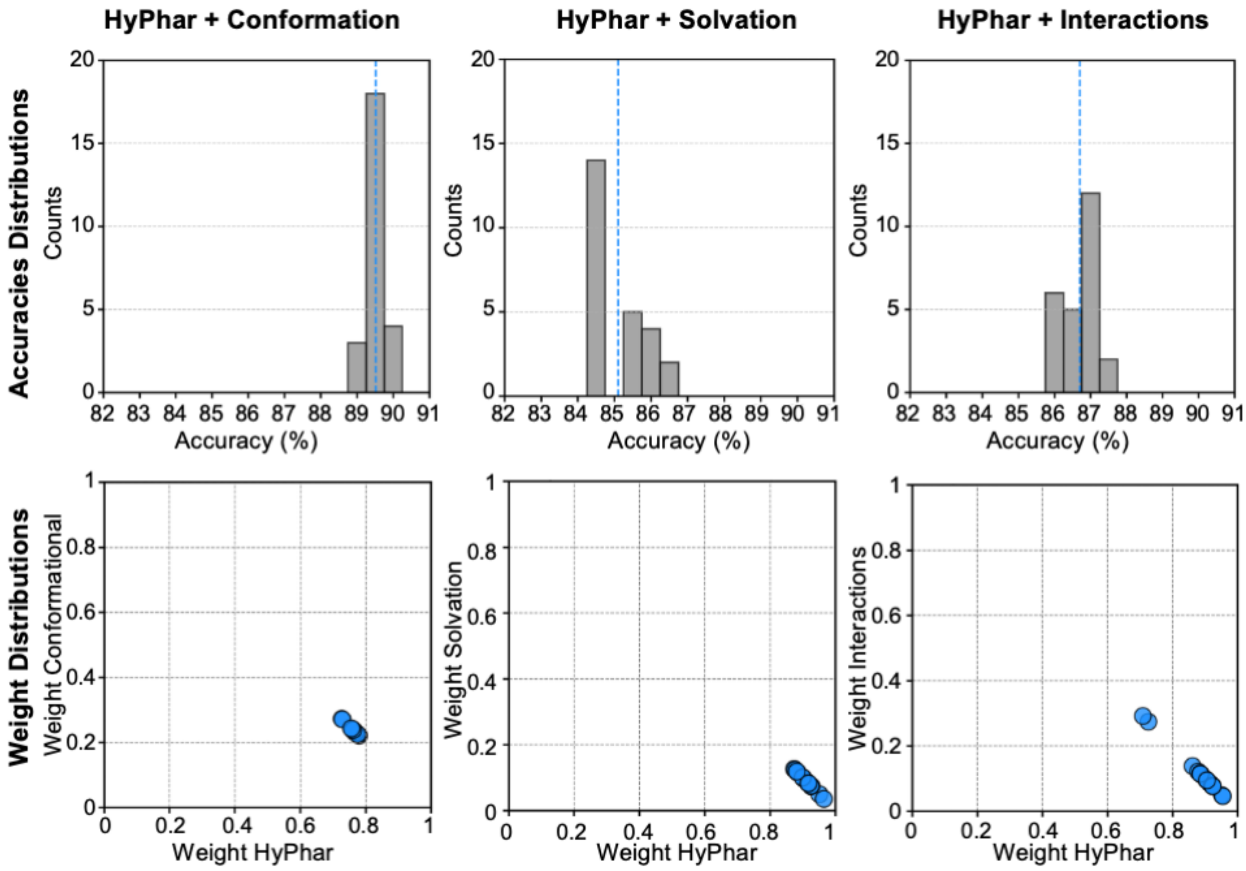


**Fig. S5. Effect of combining selected properties (conformational stress, desolvation penalty, number of interactions) on the accuracy of *S<sub>HYDRO</sub>*.** (Top) Distribution of the accuracy obtained from 25 Bayesian optimization runs combining the hydropathic score with the conformational penalty, the desolvation cost and the number of interactions determined for the ligand (X-ray, native-like and decoy) poses. (Bottom) Representation of the weights given to the hydropathic score and the conformational penalty/desolvation cost/number of interactions in the distinct Bayesian optimization runs. The weights were determined for a subset of 800 complexes taken from the DB1000 dataset, and the accuracy was estimated upon application of the weighted properties to the subset of 200 complexes included in the test set.

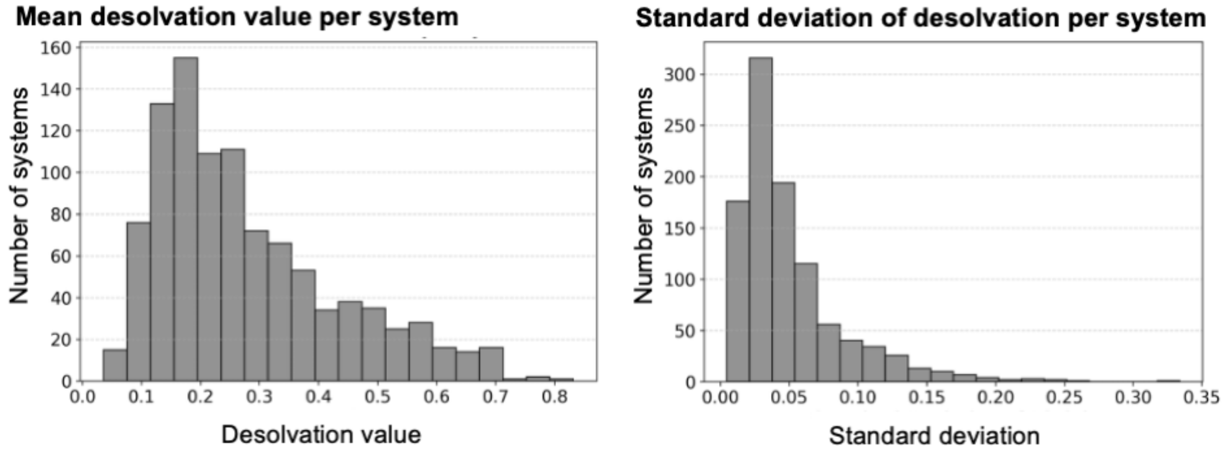


**Fig. S6. Distribution of the difference in desolvation penalty for the complexes in DB1000**. For each of the 1000 ligand-protein complexes in DB1000, the desolvation penalty was calculated for the X-ray and docked (poses 1-5) ligand using Eq. 10. The plots show the distribution of (left) the average values of desolvation penalty and (right) the standard deviation determined for both X-ray and docked poses.

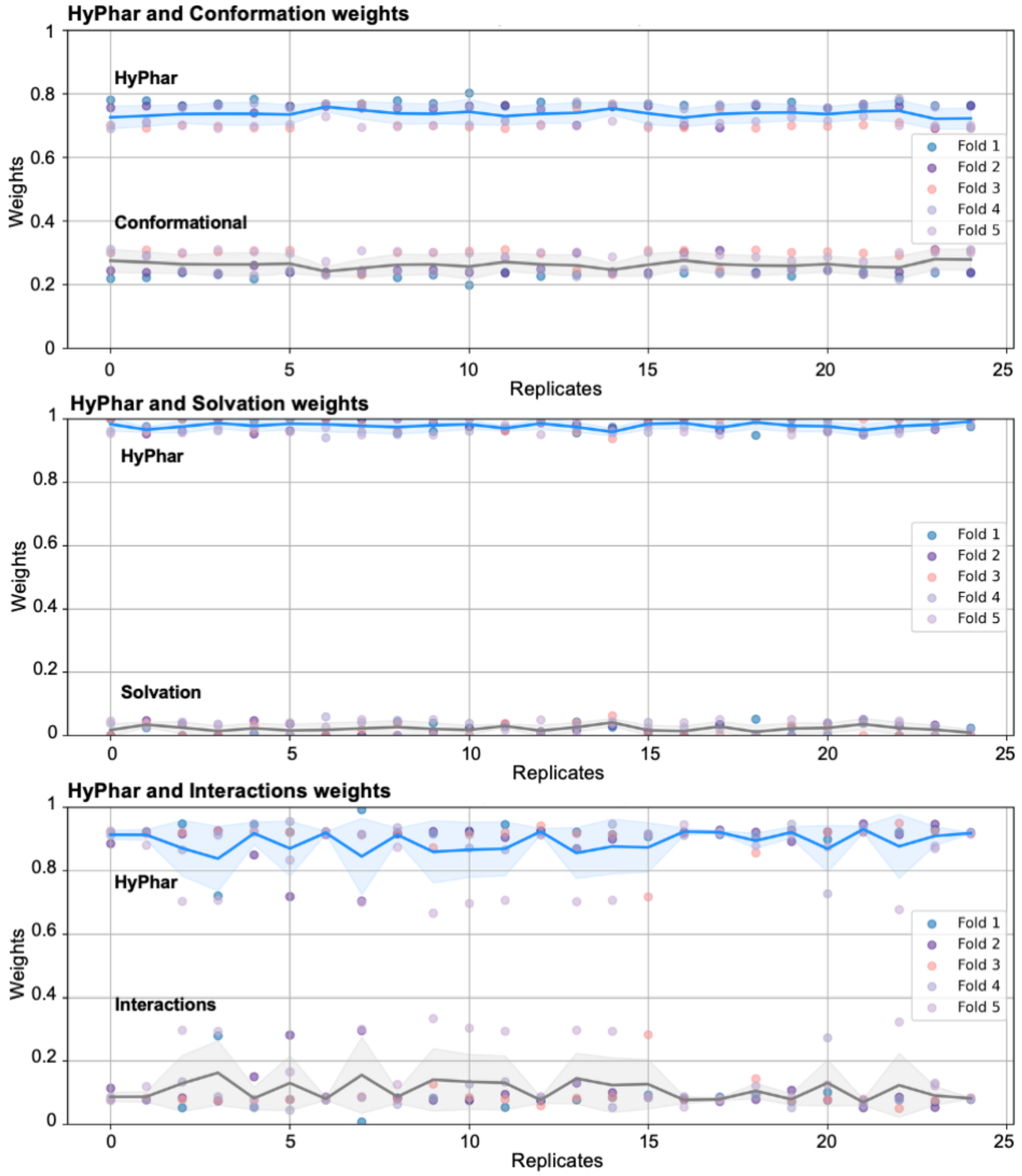


**Fig. S7. Cross-validation of the weights determined for hydropathic score, conformational penalty, (de)solvation cost and number of interactions.** Each plot represents the distribution of weights obtained from 25 Bayesian optimization runs (replicates) performed for a common subset of 800 complexes used as training dataset. Cross-validation was performed considering 5 distinct subsets, each consisting of 800 randomly chose complexes taken from the DB1000 dataset.
